# Identification of a sensory neuron Cav2.3 inhibitor within a new superfamily of macro-conotoxins

**DOI:** 10.1101/2022.07.04.498665

**Authors:** Celeste M. Hackney, Paula Flórez Salcedo, Emilie Mueller, Thomas Lund Koch, Lau D. Kjelgaard, Maren Watkins, Linda Grønborg Zachariassen, Pernille Sønderby Tuelund, Jeffrey R. McArthur, David J. Adams, Anders S. Kristensen, Baldomero Olivera, Rocio K. Finol-Urdaneta, Helena Safavi-Hemami, Jens Preben Morth, Lars Ellgaard

## Abstract

Animal venom peptides represent valuable compounds for biomedical exploration. The venoms of marine cone snails constitute a particularly rich source of peptide toxins, known as conotoxins. Here, we identify the sequence of an unusually large conotoxin, Mu8.1, that defines a new class of conotoxins evolutionarily related to the well-known con-ikot-ikots and two additional conotoxin classes not previously described. The crystal structure of recombinant Mu8.1 displays a saposin-like fold and shows structural similarity with con-ikot-ikot. Functional studies demonstrate that Mu8.1 curtails calcium influx in defined classes of murine somatosensory dorsal root ganglion (DRG) neurons. When tested on a variety of voltage-gated ion channels, Mu8.1 preferentially inhibited the R-type (Cav2.3) calcium channel. Ca^2+^ signals from Mu8.1-sensitive DRG neurons were also inhibited by SNX-482, a known spider peptide modulator of Cav2.3 and voltage-gated K^+^ (Kv4) channels. Our findings highlight the potential of Mu8.1 as a molecular tool to identify and study neuronal subclasses expressing Cav2.3. Importantly, this multidisciplinary study demonstrates the feasibility of large, disulfide-rich venom-component investigation, an endeavor that will lead to the discovery of novel structures and functions in the previously underexplored group of macro-conotoxins.

## Introduction

Animal venom peptides and proteins are employed for the incapacitation of prey or the defense against predators and competitors (1). Venom components function by binding with high affinity and selectivity to their molecular targets. These are often specific membrane-bound proteins that control vital cellular signaling pathways and include ligand and voltage-gated ion channels, G protein-coupled receptors, tyrosine kinase receptors, and transporters (2). Because of the high similarity between venom peptide targets in the prey and their orthologs in mammals, as well as the conservation of signaling pathways, venom components often show activity in mammalian systems. Animal toxins are therefore interesting for the development of molecular probes and biological tools as well as potential drug leads. Currently, eight venom-derived drugs have been approved by the U.S. Food and Drug Administration (FDA) for human use, and approximately 30 other venom-derived peptides are in clinical and preclinical trials to treat a variety of diseases, such as diabetes, hypertension, chronic pain, thrombosis, cancer, and multiple sclerosis (3, 4).

The venom produced by predatory marine cone snails is particularly rich in peptide toxins (known as conotoxins or conopeptides). Each of the approximately 1000 extant cone snail species expresses a unique set of several hundred conotoxins (5), resulting in an estimated diversity of more than 200,000 conotoxins. Conotoxins often display exquisite specificity for their targets and are consequently used widely as pharmacological tools for research purposes. Moreover, the *ω*-MVIIA conotoxin, which inhibits the Cav2.2 calcium channel (6), is an FDA-approved drug (commercial name Prialt*®*) for the treatment of severe chronic pain (7, 8).

Conotoxins are used extensively to investigate ion channel function (and dysfunction) as illustrated by the κM-conotoxin RIIIJ from *Conus radiatus* that displays subtype selectivity for asymmetric heteromeric voltage-gated K^+^ (Kv1) channels (9, 10). The selectivity of this toxin has recently been employed to identify a new subclass of peptidergic nociceptors – sensory neurons that respond to stimuli and transmit a signal to the central nervous system (CNS) – with distinct properties (11). Somatosensory neurons comprise a heterogeneous population of neurons that can be divided into subclasses using constellation pharmacology (12). Using this approach, individual neuronal cells in a population of mouse dorsal root ganglion (DRG) neurons are screened with a combination of calcium imaging and pharmacological compounds that each elicit a characteristic response used to differentiate neuronal cell types.

Conotoxins are produced and folded in the endoplasmic reticulum (ER) of cone snail venom glandular cells. Thus, conotoxin preproproteins typically comprise a signal sequence for entry into the ER, a propeptide region of largely unknown function, and the mature peptide that is proteolytically released from the propeptide (13). Conotoxins are classified into gene superfamilies based on N-terminal signal sequence similarity (14), with more than 50 gene superfamilies identified to date (13). Some superfamilies comprise several subfamilies (which we term “classes” in the current work). For instance, this is the case for the C-superfamily that comprises the consomatin and contulakin-G classes (15). In contrast to the signal sequence, the mature peptide region exhibits remarkable sequence variability except for the presence of conserved cysteines that form disulfide bonds critical for stability. In addition to disulfide bonds, conotoxins can acquire a variety of other post-translational modifications, such as C-terminal amidation, O-glycosylation, hydroxylation, and bromination that can, in some cases, influence target binding (16–19).

The advent of new sequencing technologies and bioinformatics tools for transcriptome analysis has revealed thousands of previously unknown animal venom peptide and protein sequences in recent years (20). Although cone snails mostly express short peptide toxins (mean length of the mature peptide: 42 residues (13)), the many new available sequences reveal that they also produce larger toxins. These more complex molecules have not been comprehensively explored, mostly because of limitations in the production of large, cysteine-rich proteins. Specifically, unlike the short peptide toxins, the larger toxins are rarely amenable to chemical synthesis and subsequent *in vitro* folding. Here, we coin the term “macro-conotoxin” for this group of conotoxins generally longer than 50 amino acid residues.

In this study, we identify and investigate a previously uncharacterized conotoxin, Mu8.1, from the fish-hunting snail *Conus mucronatus.* We produce this unusually large conotoxin of 89 residues using a modified *Escherichia coli* expression system and uncover that it belongs to a distinct class of conotoxins evolutionarily related to the con-ikot-ikots as well as two hitherto unknown conotoxin classes. We demonstrate that Mu8.1 inhibits depolarization-induced Ca^2+^ influx in mouse peptidergic nociceptors, likely through targeting the R-type (Cav2.3) voltage-gated Ca^2+^ channels. In addition to identifying a new conotoxin superfamily and providing structural insight at the atomic level, this study establishes Mu8.1 as a new molecular tool for the investigation of the role of Cav2.3-mediated currents in sensory neurons. Importantly, our work also demonstrates that an understudied pool of macro-conotoxins, such as Mu8.1, is amenable to detailed structural and functional investigation.

## Results

### Mu8.1 defines a new class of conotoxins evolutionarily related to the con-ikot-ikots

Transcriptome analysis of the venom gland of *C. mucronatus,* a fish-hunting species from the *Phasmoconus* clade (20), revealed the presence of two previously unrecognized toxin transcripts. Their high degree of sequence similarity designated them to be allelic variants that we assigned the names Mu8.1 and Mu8.1ii (21). The protein sequence of Mu8.1 displays the characteristic tripartite organization of conotoxins consisting of a predicted N-terminal signal sequence (21 residues), an intervening propeptide region (5 residues), and lastly the toxin-encoding region (89 residues) (Fig. 1A). The Mu8.1 and Mu8.1ii sequences differed only in the substitution of three residues in the C-terminal region (Fig. 1B).

**Figure 1.**
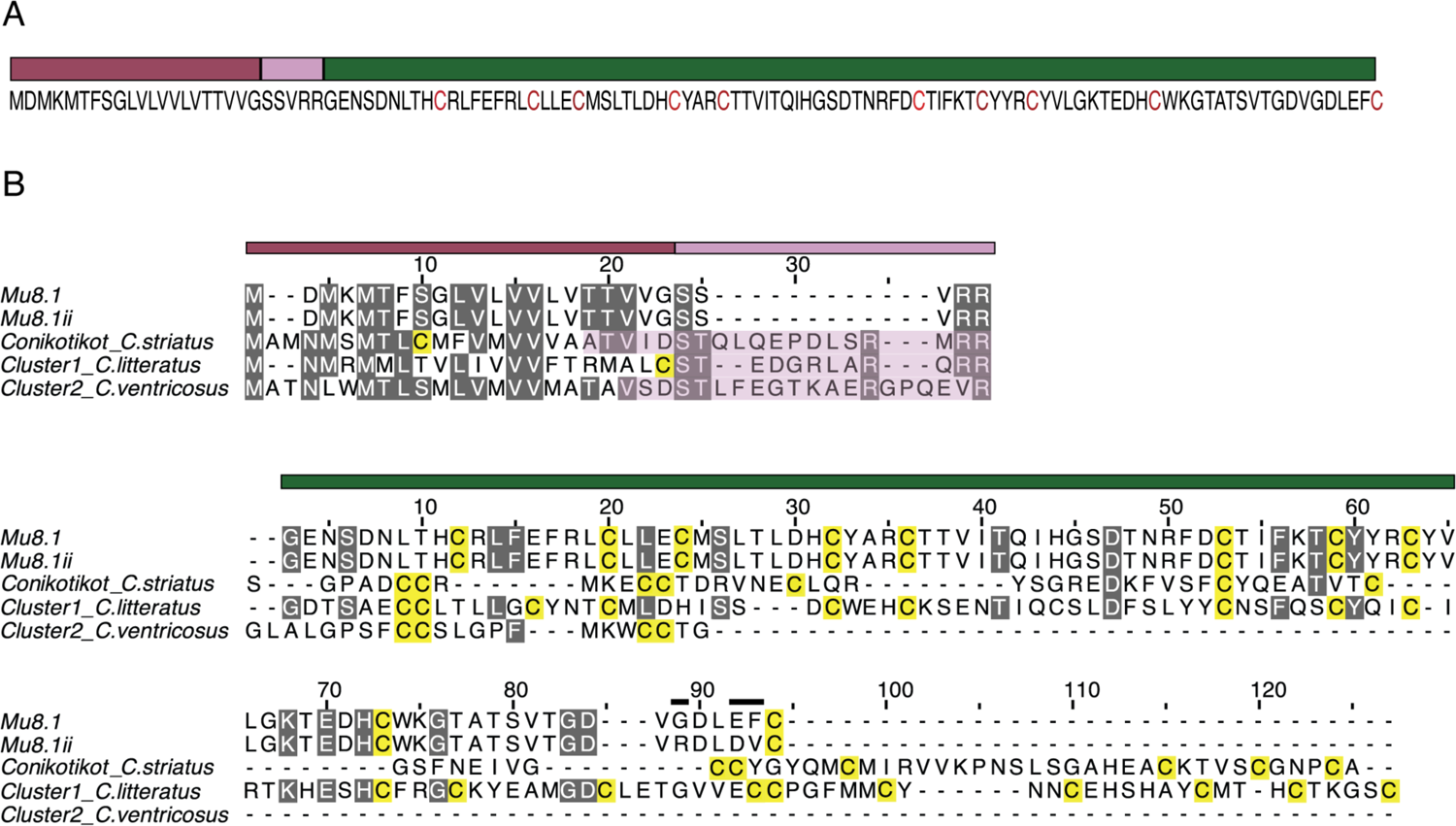
Mu8.1 belongs to a new conotoxin superfamily. **A.** Mu8.1 prepro-sequence annotated with colored bars indicating the predicted tripartite organization. Mauve: signal sequence, pink: propeptide region, green: mature Mu8.1. Cysteine residues are colored red. **B.** Alignment of Mu8.1/ii and select sequences from the con-ikot-ikot (from *C. striatus* (cs)), Cluster 1 (from *C. literatus*), and Cluster 2 (from *C. ventricosus*) classes. Top: multiple sequence alignment of the predicted signal and propeptide sequences (mauve and pink bars denote signal and propeptide organization for Mu8.1, shaded pink box portion indicates the propeptide sequences of con-ikot-ikot, Cluster 1 and Cluster 2). Bottom: alignment of the mature toxin sequences (green bar). The three sequence differences between Mu8.1 and Mu8.1ii are marked with black bars above the residues in the alignment. Gray: columns with at least 60% sequence identity, yellow: cysteine residues.

To uncover Mu8.1-related sequences from other *Conus* species, we performed a BLASTp search (22) using the Mu8.1 precursor as the query sequence. Several of the identified sequences were annotated as con-ikot-ikot toxins, including sequences from three worm-hunting species, *Conus praecellens* (23), *Conus andremenezi* (23), and *Conus vexillum*. Three of the toxins from *C. vexillum*, Vx65, Vx78, and Vx79, have been identified in venom extracts by MS/MS (24). However, con-ikot-ikot signal sequences are different from the signal sequences of Mu8.1 and Mu8.1ii (Fig. 1B) and those observed in the sequences from the worm-hunting species mentioned above (Fig. S1), contradicting the database annotation of these peptides as con-ikot-ikot toxins. Moreover, the arrangement of cysteines in the mature sequence of Mu8.1 and homologous toxins is also distinct from what is observed in con-ikot-ikot toxins (see further below).

To identify conotoxins belonging to the same class as Mu8.1, we mined the transcriptome data available in the NCBI, DDBJ, and CNGB sequence repositories. Searching a total of 602,377 assembled transcripts from these venom gland datasets (Table S1), we identified an additional 104 sequences from 38 species. These sequences included examples from the snail-hunting species *Conus marmoreus*, *Conus textile,* and *Conus gloriamaris*, as well as several from various worm-hunting species. The identified signal sequences are highly conserved and harbor a MDMKMTFSGFVLVVLVTTVVG consensus sequence (Fig. S1). Several of the sequences contain two additional N-terminal residues, MT. All sequences contain 10 conserved cysteine residues in the region of the mature toxin with identical loop sizes between them, with a few sequences containing one or two additional cysteines. As is common in conotoxins, there is a high variability between the mature sequences. Still, 10 residues (in addition to the cysteines) are strictly conserved among species (Fig. S1). Based on the structural similarity with proteins carrying a saposin domain (see further below) we refer to this new class of toxins as saposin-like conotoxins (SLCs).

The database annotation of several SLCs as con-ikot-ikots led us to investigate a potential distant evolutionary relationship between these two classes. To do so, we identified all published *Conus* sequences with sequence similarity to con-ikot-ikots and grouped these using CLANS clustering analysis. This method uses all-against-all BLAST e-values to attract sequences with high similarity and repel sequences of little similarity, thereby forming clusters of highly similar sequences. The CLANS analysis (Fig. S2) showed four distinct but interconnected clusters corresponding to the identified toxin classes: the ‘classical’ con-ikot-ikots, the SLCs, and two previously unrecognized conotoxin classes (Clusters 1 and 2). The numerous links interconnecting the four clusters suggest a common evolutionary origin. To further probe the putative evolutionary relationship between the sequences of the four classes we compared the nucleotide sequences of their 5’UTRs and the beginning of their ORFs using randomly selected sequences from each class. This sequence alignment showed a high sequence identity of the 5’ UTR in the four toxin classes (Fig. S3), further suggesting a common evolutionary origin. The conclusion that the sequences of four identified classes are related through an ancestral gene was corroborated by the finding that their gene structures are very similar, with all introns occurring in the same phase (phase 1) (Fig. S4). We concluded that the four identified classes – SLC, con-ikot-ikot, Cluster 1, and Cluster 2 – together comprise a new superfamily where the individual classes have related signal sequences, but distinct mature regions (Fig. 1B).

### Recombinantly expressed Mu8.1 purifies as a single, fully oxidized species from *E. coli*

To gain insight into the structure and function of the new SLC family of toxins, we expressed Mu8.1 using the csCyDisCo system (25). This system, based on the original CyDisCo system (26, 27), allows disulfide-bond formation in the cytosol of *E. coli* as a result of expression of the Erv1p oxidase along with two protein disulfide isomerases (human PDI and a conotoxin-specific PDI) from an auxiliary plasmid (25, 28). Mu8.1 was expressed as a fusion protein with an N-terminal ubiquitin (Ub) tag containing 10 consecutive histidine residues inserted into a loop region followed by a TEV-protease recognition site (Fig. S5A). The beneficial effect of the csCyDisCo system was confirmed by SDS-PAGE gel and Western blot analysis, where the Ub-Hi_10_-Mu8.1 fusion protein was found in the soluble fraction (to approximately 50%) only when co-expressed with Erv1p and the two PDIs (Fig. 2A), as we have previously observed for other conotoxins (25, 28). TEV protease cleavage of Ub-Hi_10_-Mu8.1 resulted in the liberation of the 89-residue mature conotoxin, which was purified (>95% purity; Fig. 2B) in a final yield of *∼*1 mg per liter of culture.

**Figure 2.**
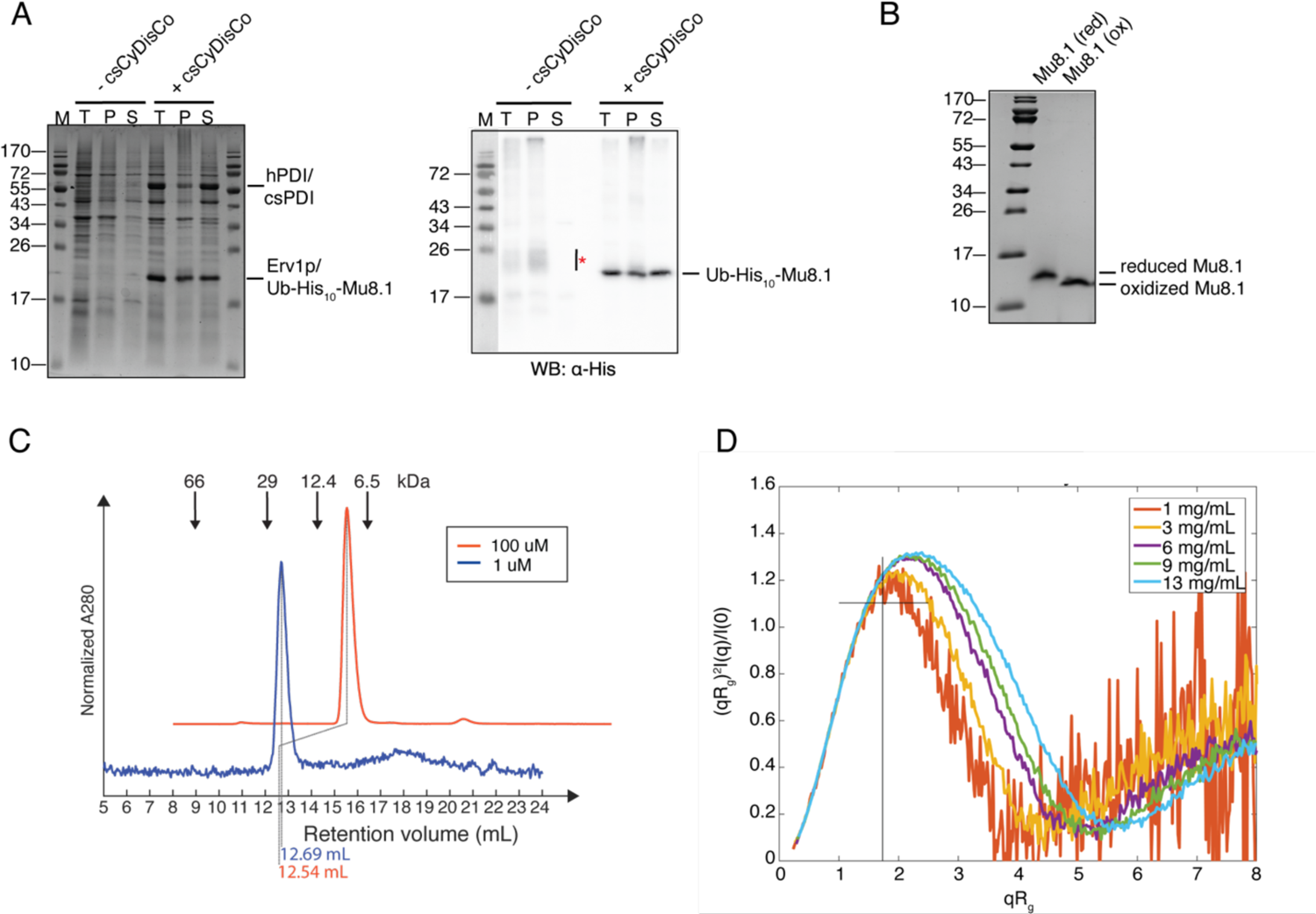
Recombinantly expressed Mu8.1 exists primarily as a non-covalent dimer in solution. **A.** Comparison of Ub-His_10_-Mu8.1 expressed in the absence (-csCyDisCo) or presence (csCyDisCo) of the csCyDisCo expression system in *E. coli* BL21(DE3) cells. Left panel: 15% SDS-PAGE gel stained with Coomassie Brilliant Blue showing the total cell extract (T), resuspended pellet after lysis and centrifugation (P), and the soluble fraction (S) from cells expressing Ub–His_10_–Mu8.1. Note that Ub-His_10_-Mu8.1 and Erv1p migrate similarly. Protein levels are directly comparable between lanes. Right panel: western blot probed with an anti-His antibody (*α*-His) for detection of Ub-His_10_-Mu8.1 in the same samples loaded on the SDS-PAGE gel. **B.** 15% Tris-Tricine SDS-PAGE gel analysis of purified Mu8.1 in the reduced (40 mM DTT; red) and oxidized (ox) state. **C.** Analytic gel filtration of Mu8.1 at 100 µM (orange) and 1 µM (blue) concentrations. Arrows denote the elution volumes of standard proteins relating to the blue trace. The dotted lines show the retention volumes of Mu8.1 in each sample. The peak intensity of the two samples was min-max normalized to allow comparison and the orange plot was offset for clarity. **D.** Normalized Kratky plot from small angle X-ray scattering analysis of five concentrations of purified Mu8.1 (1 mg/ml, orange; 3 mg/ml, yellow; 6 mg/mL, purple; 8 mg/mL, green; 13 mg/mL, light blue). The cross denotes the peak position of a globular protein.

Disulfide-bond formation in Mu8.1 was verified by SDS-PAGE analysis and full oxidation by MALDI-TOF mass spectrometry. Upon reduction, a clear mobility shift was observed by SDS-PAGE (Fig. 2B), and MALDI-TOF mass spectrometry of RP-HPLC-purified Mu8.1 verified the presence of a single species with a mass of 10181.7 Da (Fig. S5B), a value that fits the theoretical average mass of fully oxidized Mu8.1 of 10181.5 Da.

### Mu8.1 is an *α*-helical, dimeric protein in solution

More detailed insight into the structural properties of Mu8.1 in solution was obtained using CD spectroscopy, analytical gel filtration, and SAXS measurements.

The CD spectrum of Mu8.1 showed characteristic features of an *α*-helical structure with minima at 208 nm and 218 nm, and a global maximum at 195 nm (Fig. S5C). To probe the quaternary structure of Mu8.1 in solution, we first performed analytical size exclusion chromatography on Mu8.1. Regardless of the concentration (1 µM or 100 µM), Mu8.1 eluted as a single, symmetric peak with an apparent molecular weight of 22-24 kDa, suggesting a dimer under these conditions (Fig. 2C).

To corroborate this result, SAXS measurements were carried out at five protein concentrations ranging from 1 mg/mL to 13.2 mg/mL (100 µM to 1.3 mM). Here, at low q-values, we observed an increase in the intensity with increasing protein concentration, indicating an increase in size (Fig. S6A, Table S2). A dimensionless Kratky plot of the data revealed a shift in peak position with increasing concentration, suggesting a change from a spherical conformation concomitant with increasing concentration (Fig. 2D). Singular value decomposition was employed to assess the number of species present, followed by calculation of their relative volume fractions (Table S3). The results revealed the presence of three oligomeric species, with the dominant being a dimer (85% at 1 mg/mL) and increasing occurrence of tetrameric and hexameric species that correlated with increasing concentration. No monomer was observed.

Based on evidence from the analytic gel filtration and SAXS measurements, we concluded that Mu8.1 exists primarily as a dimer in solution.

### The crystal structure of Mu8.1

To elucidate key structural features of the SLC superfamily, we determined crystal structures of Mu8.1 from two different crystal conditions referred to as Mu8.1_38 and 59 (see Materials and Methods) at resolutions of 2.33 Å and 2.1 Å, respectively (Fig. 3). The asymmetric unit of Mu8.1_59 accommodates six molecules that form three equivalent dimers (Fig. S7). The asymmetric unit of Mu8.1_38 includes two molecules that form a single dimer with an interface equivalent to the one present in the three dimers in Mu8.1_59. Each protomer of the Mu8.1 structure comprises two regions (Fig. 3A and B). The first region comprises an N-terminal 3_10_-helix (3_10_N), followed by two α-helices (α1 and α2), and a 3_10_-linker helix (3_10_L). The second region comprises two α-helices (α3 and α4), and the C-terminal 3_10_-helix (3_10_C). The disulfide bonds are found primarily within each of these two regions. Within the first region, the Cys18– Cys34 and Cys22–Cys30 disulfide bonds connect α1 and α2. Further, the 3_10_N-helix connects to α3 through the Cys10–Cys51 disulfide bond. In the second region, Cys61–Cys71 links α3 and α4, and Cys57–Cys89 tethers the C-terminus to α3 (Fig. 3B).

**Figure 3.**
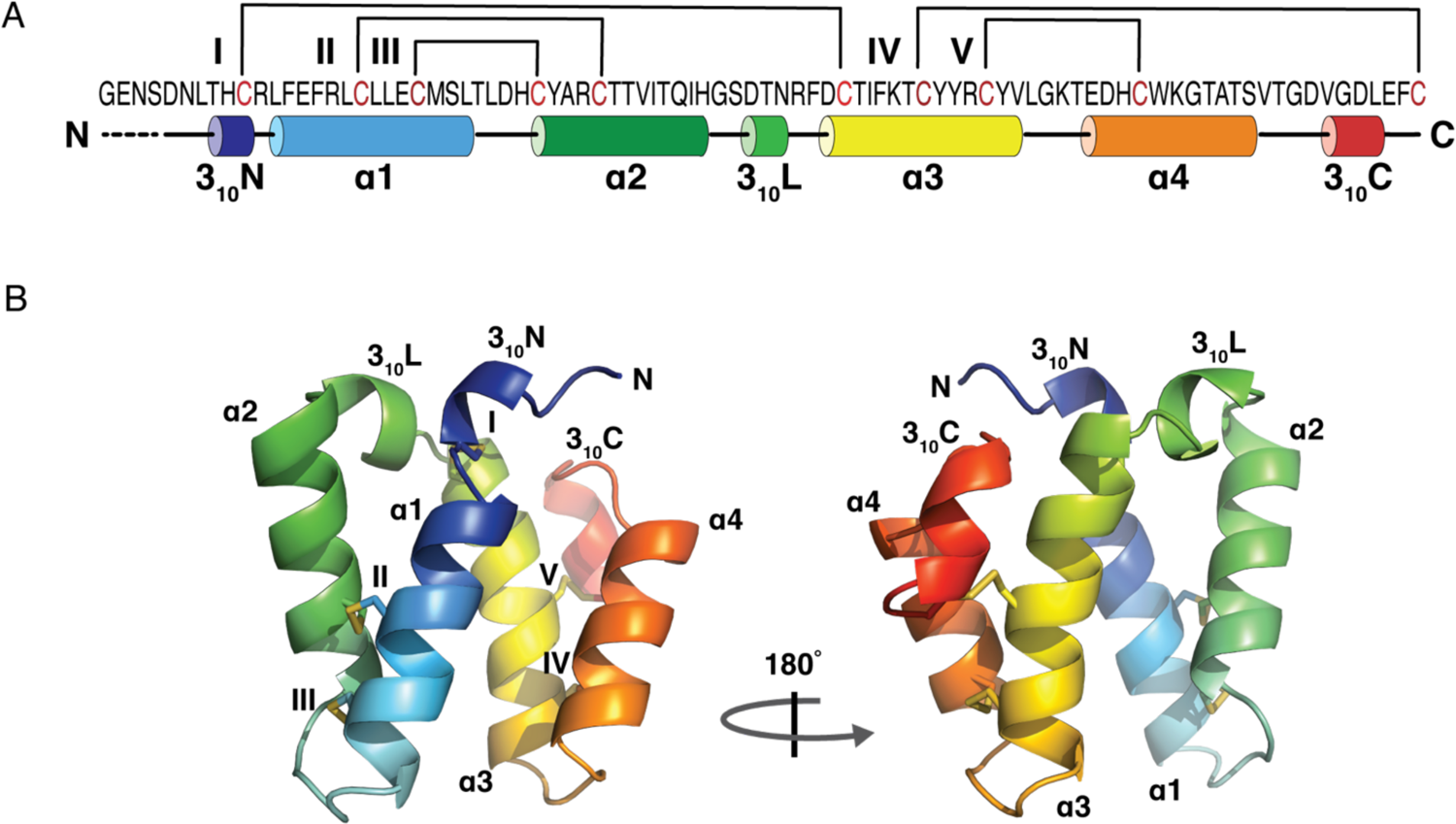
Crystal structure of Mu8.1. **A.** Graphic representation of the Mu8.1 structure with disulfide bridges represented by brackets (numbered by roman numerals) and *α*- and 3_10_-helices represented by cylinders (numbered by arabic numerals). **B.** Cartoon representation of the Mu8.1 protomer containing three 3_10_ helices interspersed with four *α*-helices labelled as *α*1-*α*4. The disulfide bridges are shown as yellow stick models and labelled with roman numerals as in Panel A.

The two helical regions surround a hydrophobic core predominantly formed by aromatic residues (Phe15, Tyr31, Tyr58, Tyr59, Trp72; Fig. 4A). Moreover, α1, α3, and α4 are held together by a network of contacts between Arg16, Tyr58, Glu68, and Trp72, where ionic interactions are present between the side chains of Arg16 and Glu68, whereas Tyr58 and Trp72 partake in T-shaped pi-pi interactions (Fig. 4A). This network of residues is completely or highly conserved among the SLC superfamily toxins (Fig. S1). The only variations are seen in four sequences where Arg16 is substituted with a Lys and Tyr58 is substituted by a Phe.

**Figure 4.**
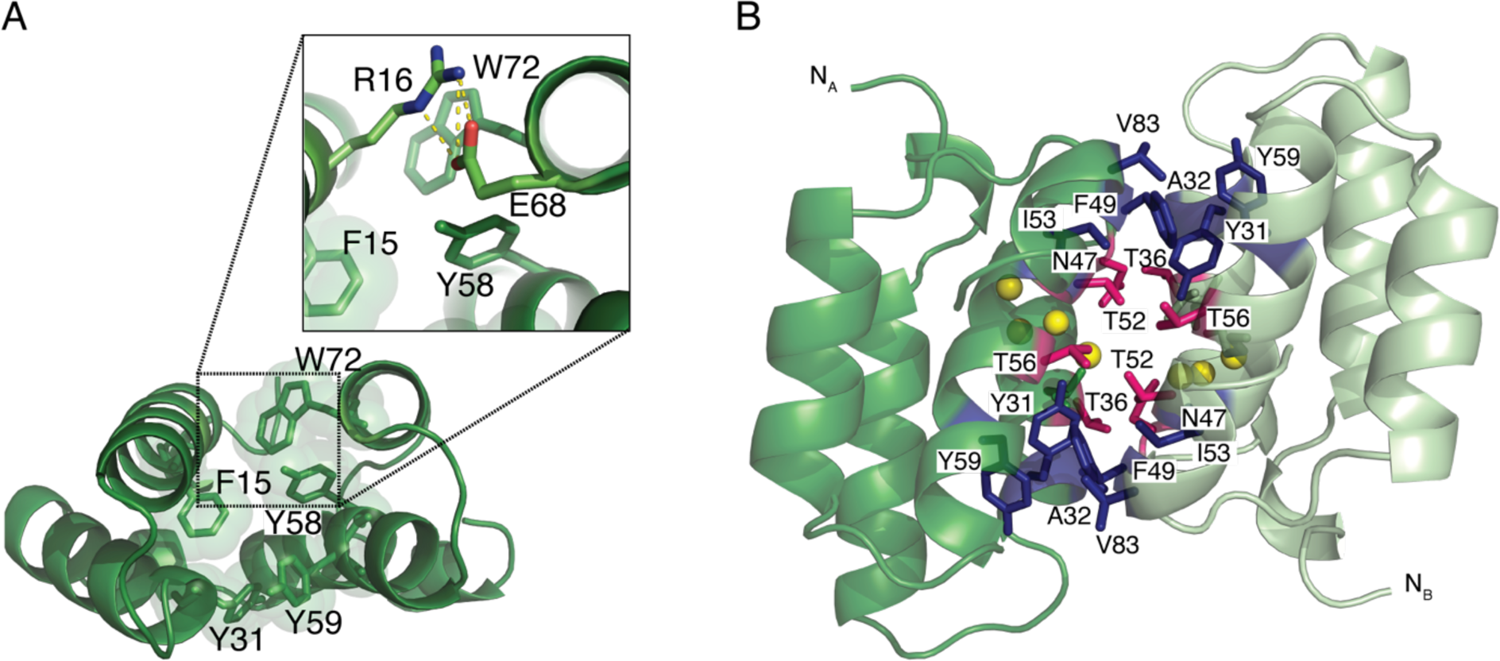
Crystal structure of dimeric Mu8.1. **A.** Cartoon representation of the Mu8.1 protomer, with the aromatic residues (F15, Y31, Y58, Y59, and W72) contributing to the hydrophobic core depicted as stick models and the corresponding van der Waals radii depicted as translucent spheres. Zoom: Select conserved amino acid residues – R16, Y58, E68, and W72 (shown as stick models) – whose interactions play a structural role (see main text for details). Ionic interactions are shown as yellow dotted lines. **B.** “Side view” of the Mu8.1 dimer highlighting the amino acid residues (depicted as sticks) composing the dimer interface. Protomer A is shown in forest green, and protomer B shown in pale green. Hydrophobic residues are in blue and polar residues in pink. Water molecules are represented by yellow spheres.

The dimer interface is characterized by both hydrophobicity as well as the presence of a water-filled cavity surrounded by the polar residues Thr36, Asn47, Thr52, and Thr56 (Fig. 4B). The residues forming the largely dry (water-excluded) hydrophobic interface include Tyr31, Ala32, Phe49, Ile53, Tyr59, and Val83. While the residues involved in structural stabilization of the Mu8.1 monomer are highly conserved among SLC superfamily proteins, the amino acid residues comprising the dimer interface exhibit a higher degree of variability (Fig. S1). A potential functional role of four additional conserved residues without any apparent structural purpose – Lys55, Arg60, Lys66, and His70 – remains speculative at present but could represent residues that interact with the molecular target of this toxin family.

### Mu8.1 has structural similarity with con-ikot-ikot but does not block GluA2 desensitization

Structural similarity often correlates with similar functional properties and, therefore, may provide important insight into protein function. We, therefore, searched the entire Protein Data Bank (PDB) for structures exhibiting similarity to Mu8.1. The search identified 48 hits showing a high degree of similarity to the Mu8.1 protomer structure, with most structures displaying a saposin-like fold (see below). Notably, con-ikot-ikot from *Conus striatus* also showed structural similarity with Mu8.1. The schematic overview of Mu8.1, con-ikot-ikot, and human saposin A (SapA) in Fig. 5A reveals the similar architecture of these proteins.

**Figure 5.**
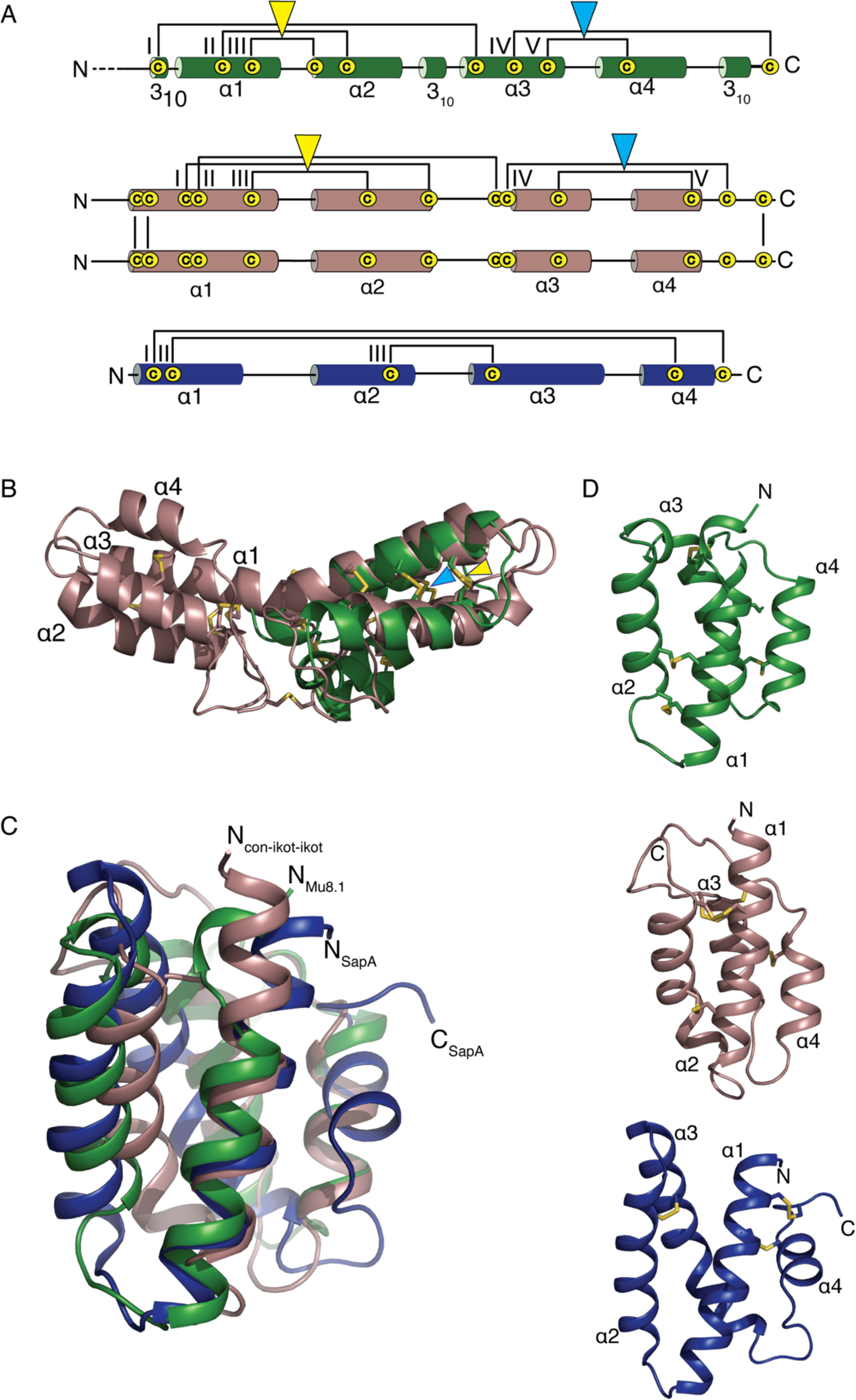
Monomeric Mu8.1 exhibits a Saposin-like fold. **A.** Graphic representations of Mu8.1 (green), con-ikot-ikot (dirty violet), and SapA (blue). Con-ikot-ikot is a homodimer held together by three intermolecular disulfide bonds, with each subunit comprising four *α*-helices and five disulfide bonds (31). SapA comprises four *α*-helices and three disulfide bonds. Disulfide bonds are depicted as brackets and labelled with Roman numerals. Amino acid residues not resolved in the crystal structure of Mu8.1 are shown as dotted lines. The yellow and light blue triangles denote structurally conserved disulfide bonds between Mu8.1 and con-ikot-ikot. **B.** Overlay of cartoon representations of monomeric Mu8.1 (forest green) and con-ikot-ikot dimer (PDB:4U5H) (dirty violet). Disulfide bonds are represented as yellow sticks. Overlay was performed using the CLICK server as described in the Materials and Methods section and visualized in Pymol. **C.** Overlay of cartoon representations of monomeric Mu8.1 (forest green), monomeric con-ikot-ikot (dirty violet), and SapA (PDB: 2DOB) (blue). Con-ikot-ikot and Mu8.1 superposition was performed with the Click server as described in Materials and Methods. SapA was manually overlaid with the two other molecules in Pymol using PyMol’s super command. **D.** Mu8.1 (green), con-ikot-ikot (dirty violet), and SapA (blue) shown separately in the same orientation as in Panel C.

Con-ikot-ikot targets the AMPA-type of ionotropic glutamate receptors and has been shown to block desensitization of the receptor producing a sustained agonist receptor-mediated current (29, 30). The crystal structure of con-ikot-ikot, determined in complex with the GluA2-type AMPA receptor, has shown a homo-dimeric protein covalently linked by disulfide bonds formed between three of the 13 cysteine residues present in each monomer (31) (Fig. 5A). Despite low sequence similarity between con-ikot-ikot and Mu8.1 (Fig. 1B), the structures of the Mu8.1 protomer and con-ikot-ikot superimpose well with an RMSD of 2.14 Å for all C*α* atoms (Fig. 5B). However, despite this high degree of structural similarity between the two proteins, Mu8.1 did not block the desensitization of the GluA2 AMPA receptor transiently transfected into HEK293 cells (Fig. S8).

### Mu8.1 and con-ikot-ikot display a saposin-like fold

Saposin-like proteins (SAPLIPs) comprise a protein family with over 200 members that perform a variety of biological functions and are found in a phylogenetically diverse group of eukaryotes. Most of these involve lipid interaction, and lead to local disordering of the lipid structure, membrane perturbation, or membrane permeabilization (32), although the fold is also associated with other functions (33). SAPLIPs display a characteristic four-helix bundle structure stabilized by a conserved pattern of three disulfide bonds (Fig. 5A, 5C, and 5D).

SapA was one of the top hits in our search, although the SapA and Mu8.1 sequences share only 21.6% identity. The SapA and Mu8.1 have similar structural topologies and superimpose with an RMSD of 1.81 Å for all C*α* atoms, with the two first *α*-helices superimposing especially well (Fig. 5C). A significant difference is constituted by the disulfide pattern in the two proteins (Fig. 5A and 5D). The functional implications of this observation suggest that Mu8.1 is unlikely to be involved in lipid binding (see Discussion). Overall, this analysis revealed a previously undiscovered structural resemblance between con-ikot-ikot and SAPLIPs that extends to Mu8.1.

### Mu8.1 is bioactive in zebrafish and mice

Having obtained detailed structural knowledge about Mu8.1, we next turned to investigating functional properties of Mu8.1 at the organismal, cellular, and molecular levels. We first evaluated the bioactivity of Mu8.1 by assessing the changes in locomotion of adult zebrafish after intramuscular injections. Before the injection, we recorded the free-swimming behavior of a fish for 20 min and calculated the distance traveled in this conditioning phase (Fig. 6A). We then used this measurement as a baseline reading and compared it with the distance traveled following injection with Mu8.1. After injection, all zebrafish (n=3) exhibited an immediate increase in movement followed by a slower on-set decrease in locomotion compared to the saline controls (n=3) (Fig. 6B). All fish had returned to normal swimming patterns and exploratory behavior 4 hours after the injection.

**Figure 6.**
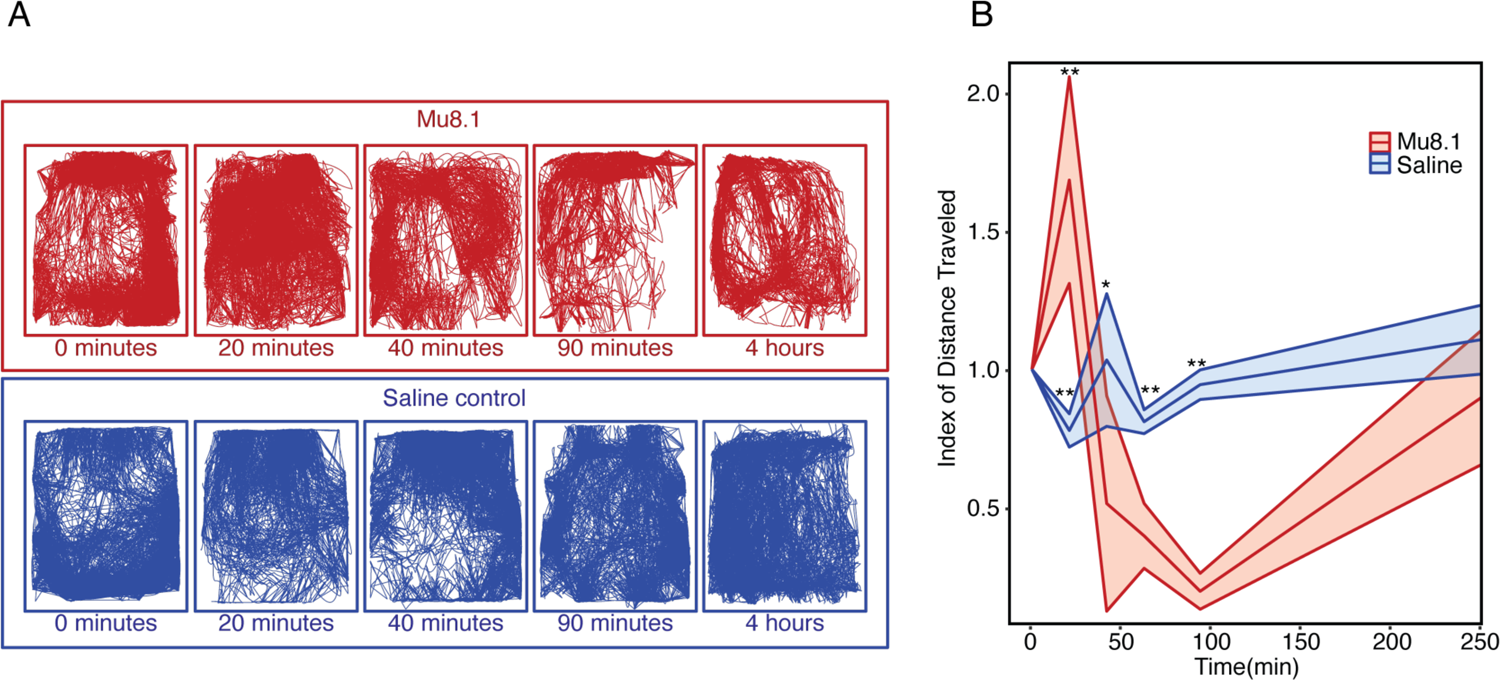
Intramuscular injection of Mu8.1 induces a behavioral response in adult zebrafish. **A.** Swimming traces after injection of Mu8.1 (5 nmol; red) or saline control (blue). Time zero represents 20 minutes of free-swimming pre-injection. **B.** Quantification of distance traveled at 20, 40, 90, and 240 minutes after intramuscular injection (n=3). The shaded regions represent standard deviation. A t-test was performed to calculate significant differences in total distance traveled between fish injected with Mu8.1 and saline solution (P ≤ 0.05= *, P ≤ 0.01 = **).

Because of their closer kinship to humans, we also analyzed the bioactivity of Mu8.1 in mice by intracranial injection as described previously (34). Ten minutes after injection, the Mu8.1 group (n=3) displayed a hypoactive phenotype characterized by reduced activity akin to a sleeping state in comparison to the vehicle control animals (n=2). The Mu8.1-treated mice were not responsive to auditory or light stimuli but reacted to a light poke to the back paws by moving away 2-3 steps. Mu8.1-induced hypoactivity persisted for ∼1 hour with a return to normal exploratory and grooming behavior occurring 2 hours post-injection in all the treated mice.

### Mu8.1 modulates depolarization-induced Ca^2+^ influxes in sensory neurons

Given the bioactivity observed at the organismal level, we next tested the effects of Mu8.1 on Ca^2+^ influx in somatosensory DRG neurons that relay sensory input to the CNS. These cells were isolated from transgenic reporter mice in which the regulatory elements of the calcitonin gene-related peptide (CGRP) drive the expression of green fluorescent protein (GFP), whereby GFP-labeling identifies peptidergic nociceptive neurons. Overall, seven major classes of sensory neurons responsible for processing the sensations of cold, heat, mechanical cues, and pain were assigned based on cell size, GFP-labeling, and isolectin B4 staining (35), as well as their responses to mustard oil (AITC), menthol, capsaicin, and conotoxin RIIIJ, according to the functional classification of somatosensory neurons proposed by Giacobassi et al. (11).

Representative examples from neurons belonging to all somatosensory subclasses assayed are shown in Fig. 7A. DRG neurons incubated in observation solution (4 mM KCl) were sequentially depolarized by pulses (15 s) of high potassium (25 mM KCl) extracellular solution, which caused stereotypical rises in intracellular Ca^2+^ concentration evidenced by the increase in Fura-2 signal. Incubation with 10 µM Mu8.1 produced a decrease in subsequent Ca^2+^ peaks elicited by the high K^+^ stimulus in 20.3 ± 3.0% of the neurons analyzed (n = 2365; Fig.7B pie chart), predominantly in peptidergic nociceptors (68% of all affected cells, green bar Fig. 7B). In addition to the peptidergic nociceptors, 10 µM Mu8.1 reduced the calcium signals of DRG neurons identified as large-diameter mechanosensors (12.6% of all affected cells) and C-low threshold mechanoreceptors (C-LTMRs; 7% of all affected cells) as shown in Fig. 7B (magenta and lilac bars, respectively).

**Figure 7.**
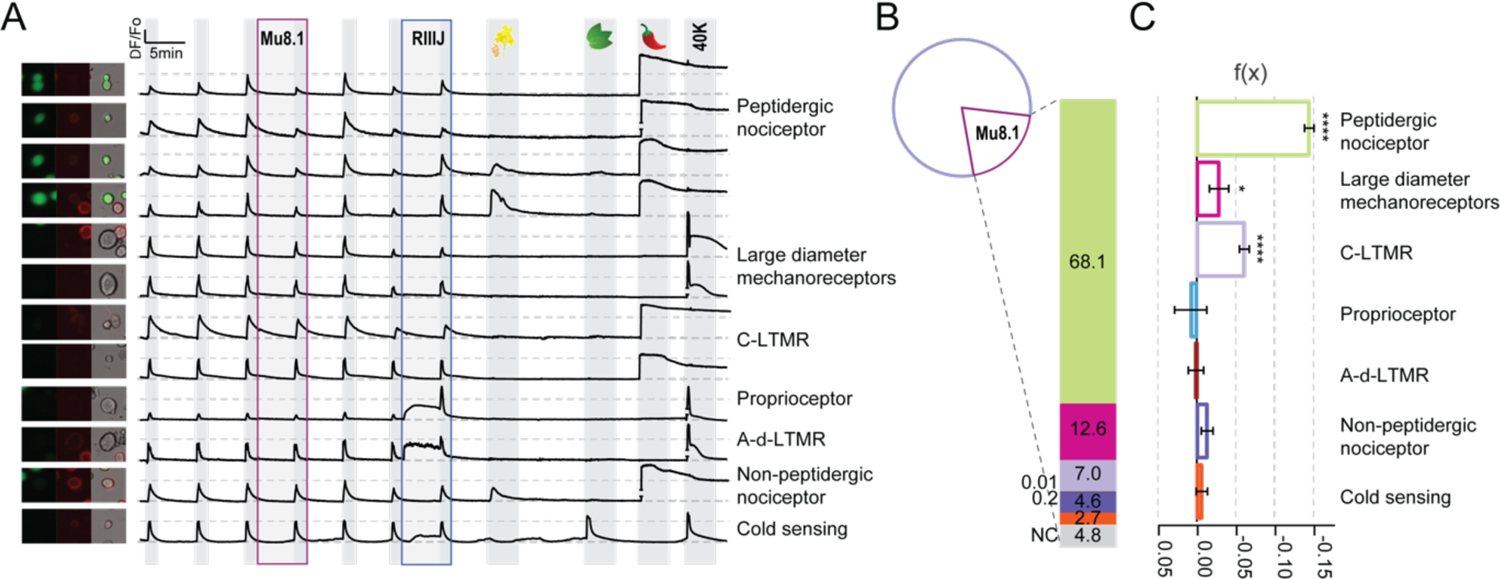
Mu8.1 inhibits calcium entry into mouse sensory neurons. **A.** Intracellular calcium changes in response to sequential pharmacological treatments from seven classes of DRG neurons (labelled on the right). Each trace represents the calcium signal (*Δ*F/Fo) of the neuron pictured on the left (GFP-CGRP+: peptidergic nociceptors; Alexa Fluor 647-Isolectin B4+: non-peptidergic nociceptors, and bright-field). KCl depolarization pulses (25 mM) are indicated by light grey shading. A higher KCl pulse (40 mM) was used to elicit maximum calcium signal at the end of the experiment. In the presence of Mu8.1 (10 µM, mauve box) KCl-induced calcium peaks are reduced in peptidergic nociceptors (traces 1-4), large-diameter mechanoreceptors (traces 5-6) and C-LTMRs (traces 7-8). Class-defining pharmacology: RIIIJ (1 µM, blue box); allyl isothiocyanate (AITC, 100 µM; mustard flower), menthol (400 µM; peppermint leaf), and capsaicin (300 nM; chili pepper). A pulse of KCl (40 mM) was used to elicit maximum calcium signal at the end of the experiment. **B.** Pie chart representing the population of neurons analyzed (n=2,365), highlighting the 20.3±3% of Mu8.1 (10 µM)-sensitive sensory neurons. The bar graph provides the percentage of each class of Mu8.1-sensitive DRG neurons. **C.** Quantification of the relative change f(x) in KCl-induced intracellular calcium signal observed in the presence of Mu8.1 (10 µM) (see Materials and Methods for normalization details). Two-tailed t-test * p ≤ 0.05; **** p < 0.0005.

Mu8.1 inhibitory effects on cytosolic Ca^2+^ concentration was estimated after Min-Max normalization (see Materials and Methods) and evaluated by two-tailed t-tests (Fig. 7C). Mu8.1 (10 μM) significantly curtailed the total influx of Ca^2+^ elicited by KCl application in peptidergic nociceptors (f(x) = –0.14 ± 0.01, n = 596; p <0.001), C-LTMRs (f(x) = –0.06 ± 0.01, n = 190; p <0.001) and Large-diameter mechanosensors (f(x) = –0.03 ± 0.01, n = 48; p = 0.0279).

### Mu8.1 inhibits recombinant and native Cav2.3-mediated currents

Given the ability of Mu8.1 to inhibit cellular Ca^2+^ influxes in DRG neurons, we assessed the modulatory effects of Mu8.1 over a comprehensive panel of recombinant voltage-gated calcium, sodium, and potassium channels (Cav, Nav, and Kv, respectively) by automated patch clamp (APC) electrophysiology as well as a large array of G protein-coupled receptors (GPCRs) by high-throughput screening assays. Among all the recombinant channels tested (Table S4), Mu8.1 was most potent against Cav2.3 channels. Fig. 8 shows representative current traces of human Cav2.3-mediated whole-cell currents exposed to increasing concentrations of Mu8.1 recorded from HEK293 cells (Fig. 8A) and the resulting concentration-response curve rendering an IC_50_ of 5.8 μM and a Hill coefficient close to unity (0.97) (Fig. 8B).

**Figure 8.**
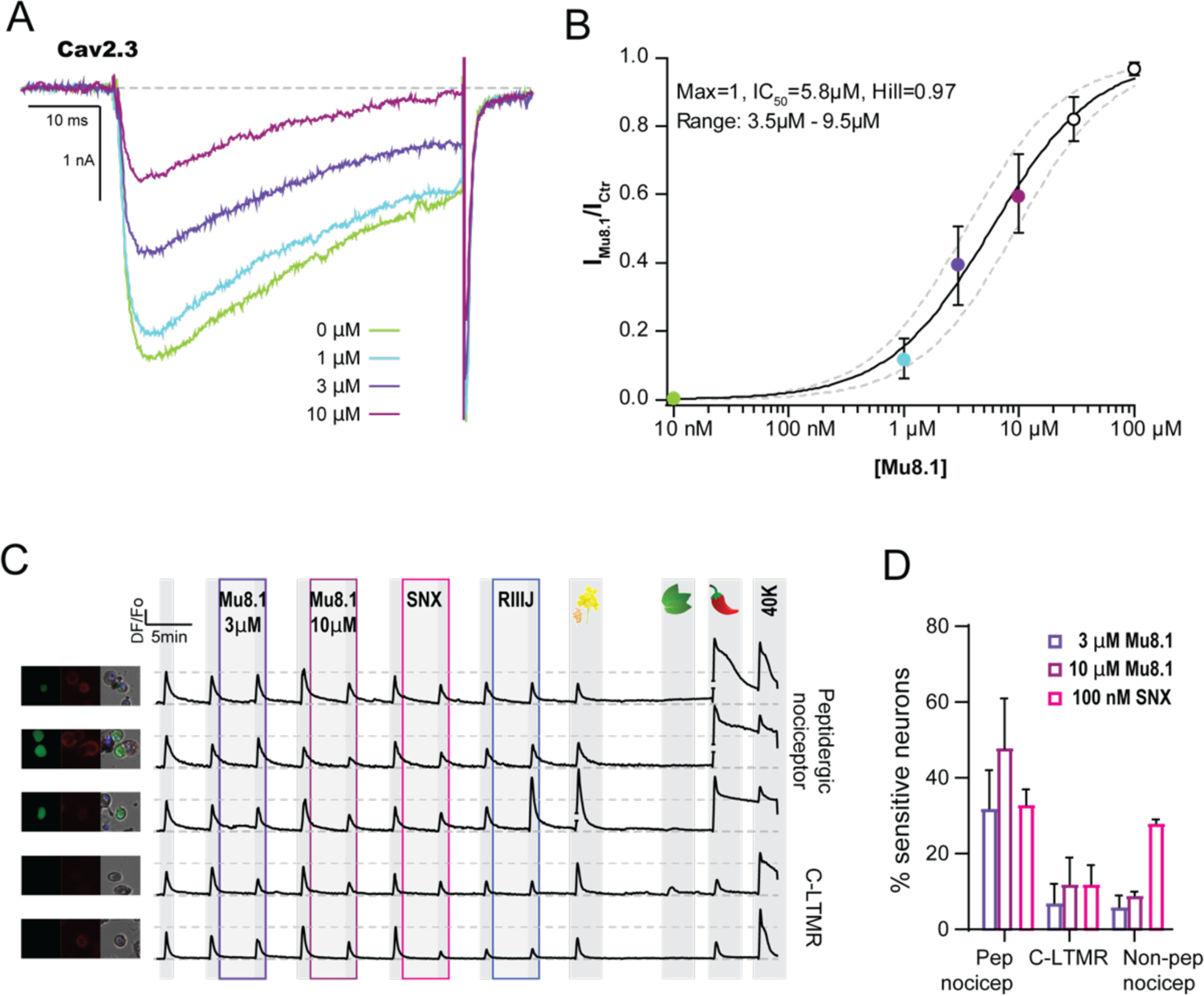
Mu8.1 inhibits recombinant human Cav2.3-mediated currents in a concentration-dependent manner. **A.** Representative whole-cell Cav2.3-mediated currents in control (green) and in the presence of 1 µM (blue), 3 µM (purple) and 10 µM (mauve) Mu8.1. Depolarization-activated currents were elicited by a 50 ms test pulse to *−*10 mV (Vh *−*80 mV; 0.1 Hz). **B.** Concentration-response relationship for the inhibition of Cav2.3 peak currents (IC_50_ = 5.8 µM; nH: 0.97; n = 5). **C.** Mu8.1 and SNX-482 target overlapping populations of peptidergic nociceptors and C-LTMRs. Cells were treated with 3 µM Mu8.1 (pink box), 10 µM Mu8.1 (mauve box) and 100 nM SNX-482 (red box). All other labels as in Fig. 7A. **D.** Relative quantification of peptidergic nociceptors (pep nocicep), C-LTMRs and non-peptidergic nociceptors (non-pep nocicep) sensitive to 3 µM and 10 µM Mu8.1 (purple and mauve, respectively) and 100 nM SNX-482 (pink). Percentages were calculated from the total number of cells recorded per class. Bars represent mean +SD from two independent DRG isolations (190-810 neurons per group).

At concentrations above 30 μM, Mu8.1 modestly inhibited Cav2.1, Cav2.2, Cav3.1-3, and Kv1.1 channels with near equipotency (IC_50_s calculated from inhibition at a single concentration of Mu8.1 are summarized in Table S4), whereas no obvious modulatory effects over heterologously expressed Nav1.4, Nav1.7, hERG, Kv1.2, Kv1.3 or Kv4.3 channels were observed (Table S4). The same was observed for a large panel of recombinant GPCRs and other receptors assessed through *β*-arrestin recruitment and radioligand binding assays (Fig. S9). Overall, our comprehensive functional screening identified the R-type Cav2.3 channel as the highest affinity mammalian target of Mu8.1, consistent with effects of Mu8.1 on DRG subpopulations, such as peptidergic nociceptors and C-LTMRs, in which Cav2.3 channels are abundantly expressed (36–38).

Using constellation pharmacology, we demonstrated that Mu8.1 inhibition of DRG neuron calcium signals were by and large reversible shortly after removal of the peptide as well as concentration-dependent. Reversibility of Mu8.1 actions can be surmised from the grey shading after Mu8.1 application in Figs. 7A and 8C, whilst 3 µM and 10 μM Mu8.1 inhibited 32 ± 10% and 48 ± 13% of subsequent calcium peaks in peptidergic nociceptors, respectively (Fig. 8C, D). Although overall reversibility was observed, 7.8% of peptidergic nociceptors did not recover after several full-bath media exchanges (Fig. S10).

SNX-482 is a spider venom peptide commonly used to study Cav2.3-mediated currents in native tissues (39) and, in contrast to Mu8.1, it is also a potent inhibitor of the Kv4 family of voltage-gated potassium channels (40). We applied SNX-482 (100 nM) to DRG neurons after Mu8.1 inhibition was reversed, verifying that these two venom-derived peptides target overlapping populations of somatosensory neurons, predominantly peptidergic nociceptors and C-LTMRs (Fig. 8C, D). Interestingly, SNX-482 also decreased Ca^2+^ signals from ∼28% of non-peptidergic nociceptors. These results strongly indicate that the decrease in Ca^2+^ influx observed upon exposure to Mu8.1 is mechanistically supported by the inhibition of Cav2.3 channels. In DRG neurons, SNX-482 not only reduced depolarization-induced Ca^2+^ signals via inhibition of Cav2.3 channels but also displayed direct and indirect Ca^2+^ signal amplification effects (Fig. S11) akin to the block of voltage-gated potassium conductances likely mediated by Kv4 potassium channel isoforms that are abundantly expressed in these neurons (41).

## Discussion

The vastly increased number of conotoxin sequences obtained in recent years from transcriptome sequencing constitutes a rich source of peptides for biomedical exploration. However, the production of new peptides often represents a bottleneck for their exploration – in particular, the characterization of large, disulfide-rich venom components is lagging. In this study, we identify the macro-conotoxin Mu8.1 from *C. mucronatus* as the founding member of the new SLC class and use a wide range of biochemical, biophysical, structural, and electrophysiological methods to provide a comprehensive characterization of the protein. Moreover, we uncover an unexpected evolutionary relationship between the SLCs and con-ikot-ikots that extends to two previously unrecognized toxin classes with all four thus defining a new conotoxin superfamily.

The relatively straightforward chemical synthesis of small peptides together with difficulties associated with the production of large venom components, in particular without *a priori* knowledge about the disulfide pattern, have biased functional studies towards small, disulfide-bridged conotoxin peptides shorter than 30 amino acid residues in length. The successful production of fully oxidized and correctly folded Mu8.1 in the csCyDisCo *E. coli* system highlights the feasibility of systematically exploring increasingly larger multi-disulfide conotoxins than was previously possible. Moreover, we recently developed the DisCoTune system based on CyDisCo to allow titration of T7 RNA polymerase repression (28). This feature permits optimization of expression conditions to potentially further increase yields by fine-tuning the expression level of the (disulfide-rich) target protein to better match the level of the helper proteins (Erv1p and PDI).

With a few notable exceptions, such as con-ikot-ikot and proteins belonging to common toxin families like conkunitzins, metalloproteases, hyaluronidases, and Phospholipase A_2_s (42–45), macro-conotoxins are generally unexplored. The large size of Mu8.1 and its modest potency against mammalian Cav2.3 channels prompts questions regarding the evolutionary advantage that producing and deploying this peptide may convey to *C. mucronatus*. In general, the size of macro-conotoxins likely confers specific properties not available to small toxins. For instance, larger toxins could participate in multivalent interactions with their targets, as noted for con-ikot-ikot and recently pointed out for bivalent venom peptides containing two homologous domains connected by an interdomain linker (46). It is conceivable that non-covalent dimers, as seen in Mu8.1, could also allow interaction with, e.g., two identical subunits of a molecular target. Even in a monomeric state, large toxins may interact with different target subunits, whereas their larger binding interface may well provide higher target specificity.

We unexpectedly found that Mu8.1 is structurally similar to con-ikot-ikot and that both display a saposin-like fold. This result raised the possibility that Mu8.1 may perform a function involving lipid interactions. Human SapA and its homologs SapB, SapC, and SapD are small, non-enzymatic proteins required to break down glycosphingolipids within the lysosome (47). In the absence of lipid, SapA adopts a characteristic monomeric, closed conformation where *α*1 and *α*4 (held together by two disulfides) form the stem, and *α*2 and *α*3 (connected by one disulfide bond) form a hairpin region (Fig. 5A and D). In the presence of lipids, SapA opens to expose a concave, hydrophobic surface for lipid binding. The primary areas of rearrangement are the loops between *α*1/*α*2 and *α*3/*α*4 that together operate as a hinge (48, 49). However, in contrast to SapA, the Mu8.1 structure is highly constricted by the network of disulfide bonds that “crosslinks” the molecule (Fig. 5A). Consequently, Mu8.1 is not likely to undergo a large conformational change to adopt an open conformation and, therefore, probably does not function in lipid binding in the same manner as the SAPLIPs.

Despite using a combination of multiple BLAST algorithms and hidden Markov models of whole toxin sequences, individual exons, introns, and 5’ and 3’ UTRs against published genomes and transcriptomes from several tissues, we did not identify a good candidate for an ancestral, endogenous gene that gave rise to the SLC sequences. Likewise, an analysis of known *Conus* proteins harboring a saposin-like domain (Fig. S12) did not provide an obvious candidate for an ancestral endogenous gene due to their limited similarity to the toxins and distinct gene structures. Based on the evolutionary analysis, we find it probable that the SLCs (and the other three toxin classes) evolved from a common ancestral gene and then diverged to a point where only the signal sequences and gene structures reveal their common ancestry. At the same time, we note that the structures of Mu8.1 and con-ikot-ikot display the same overall fold. Interestingly, AlphaFold structure predictions of Cluster 1 sequences show that these proteins are likely to also contain a saposin-like domain (Fig. S13). Therefore, we speculate that the ancestral gene from which the four classes evolved harbored a saposin-like domain, which has been retained in at least three of these classes throughout evolution.

Alternatively, the Mu8.1 and con-ikot-ikot sequences emerged as a result of convergent evolution. Thus, the saposin fold could constitute a “privileged” scaffold that has been selected during evolution due to favorable properties that confer structural stability and the ability to accommodate sequence variation. Such privileged scaffolds are found in a variety of toxin peptides and proteins that are functionally unrelated. Examples include the inhibitor cystine knot, granulin, defensin, and Kunitz folds (25,50,51).

Functionally, neither the monomeric nor dimeric states of Mu8.1 seem compatible with AMPA-receptor binding, thus rationalizing the observed lack of Mu8.1 effects on GluA2 desensitization (Fig. S8). First, the GluA2-binding surface of con-ikot-ikot corresponds to the dimer interface of Mu8.1. Second, three of the con-ikot-ikot residues shown to be important for GluA2 binding (Gln37, Glu48, and Ala86) are not conserved in Mu8.1 (Ala32, Asn47, and Phe49) (Fig. S14A). Third, the GluA2-binding surface of con-ikot-ikot is negatively charged (Fig. S14B) whereas the corresponding surface of Mu8.1 is predominantly positively charged (Fig. S14C). Fourth, although the Mu8.1 dimer surface is negatively charged as in con-ikot-ikot, the two dimers are of unequal dimensions (Fig. S14D).

The voltage-gated Cav2.3 channel was identified as the highest affinity target of Mu8.1 among an extensive collection of mammalian ion channels and GPCRs. Most of the knowledge about the function of Cav2.3 has been obtained from animal knockout and cellular knockdown experiments, where the channel was linked to epilepsy, neurodegeneration, and pain (52–55). In contrast to SNX-482, combined results from calcium-imaging and electrophysiology measurements indicate that Mu8.1 selectively modulates sensory neurons via inhibition of Cav2.3 channels without evidencing cross-actions against other neuronal conductances typically associated with somatosensory subclasses. Thus, Mu8.1 may represent a useful addition to the molecular toolbox available for the study of Cav2.3 physiological functions, as well as the role it plays in synaptic signaling and neuromodulation.

At a general level, this study illustrates that a combination of data mining and recombinant expression in *E. coli* can pave the way for a detailed analysis of structural and functional features of newly identified, macro-conotoxins. We propose that with the advent of *E. coli* expression systems such as csCyDisCo and DisCoTune (25,28,56), as well as others (57, 58), the time is ripe to begin the systematic exploration of a new realm of macro-conotoxins. These efforts will help provide a better understanding of the biological correlates of having large venom components, with the overarching aim of connecting the biochemical and molecular characteristics of venom components with the biology and behavior of cone snails.

## Materials and Methods

### Venom gland transcriptome analysis

RNA extraction and transcriptome sequencing and assembly were performed as described previously (59, 60). Assembled transcripts were annotated using a blastx search (22) (E-value setting of 1e-3) against a combined database derived from UniProt, Conoserver (61), and an in-house cone snail venom transcript library. The two toxins, SLC_Mu8.1 and SLC_Mu8.1ii, abbreviated as Mu8.1 and Mu8.1ii, were named according to (21). Here, Mu describes the two-letter species abbreviation (Mu for *C. mucronatus*), 8 describes the cysteine scaffold, and the number 1 represents the first toxin to be described from this gene family. The suffix ii is given to a sequence variant that likely represents an allelic variant.

### Transcriptome mining of the NCBI, DDBJ and CNGB databases

To find sequences that share similarity with Mu8.1 we mined the transcriptomes of 37 cone snail venom gland transcriptomes available in the NCBI, DDBJ, and CNGB repositories using the precursor sequence of Mu8.1 as query (accession numbers provided in Table S1). Transcriptome assemblies were done as described previously (59, 60). Signal and propeptide sequences of the identified homologous sequences were predicted using ProP v. 1.0 (62). Mature toxin sequences were predicted to begin after the last basic amino acid residue preceding the first cysteine in the sequence. Therefore, further trimming of sequences to meet this criterion was executed manually as needed. Multiple sequence alignments were carried out using the MAFFT version 7 multiple alignment online interface (63) and visualized in Jalview version 1.0 (64).

### Plasmid generation

The plasmid for bacterial expression of Mu8.1 was generated by uracil excision cloning, as described previously (65). Polymerase chain reaction was carried out using Phusion U Hot Start polymerase (Thermo Fisher Scientific) according to the manufacturer’s instructions. Based on the transcriptome data for Mu8.1, the sequence of the mature toxin was predicted as GENSDNLTHCRLFEFRLCLLECMSLTLDHCYARCTTVITQIHGSDTNRFDCTIFKTCYYRCYVLGKTEDHCWKGTATSVTGDVGDLEFC. A codon-optimized DNA sequence for bacterial expression was generated using the CodonOpt tool. Using this codon-optimized DNA sequence as a template, the following two primers were designed:

Mu8.1_sense: ACACGGAUCGGACACCAATCGTTTTGATTGCACAATCTTCAAGACCTGCTACTACCGGTGCTACGTTCTTGGTAAAACAGAAGACCATTGCTGGAAAGGGACGGCAACGTCAGTGACAGGTGATGTCGGAGATTTGGAATTTTGCTAAGAATTCGAGCTCCGTCGACAG;

Mu8.1_antisense: ATCCGTGUATCTGGGTAATAACTGTGGTACATCTCGCATAGCAGTGGTCTAATGTAAGCGACATACACTCCAACAAACACAGCCGGAACTCGAATAATCTACAATGGGTAAGGTTGTCTGAGTTTTCGCCCTGAAAATACAGATTCTCAC.

These primers were subsequently used to clone Mu8.1 into the pET39_Ub19 expression vector (66). The resulting plasmid encoding ubiquitin (Ub)-His_10_-tagged Mu8.1 (Ub-His_10_-Mu8.1) is referred to as pLE601. The fusion protein produced from pLE601 also contains a Tobacco Etch Virus (TEV) protease recognition site following the Ub–His_10_-tag. Primers were purchased from Integrated DNA Technologies and the sequence encoding Ub-His_10_-Mu8.1 was confirmed by Eurofins.

### Protein expression

Chemically competent *E. coli* BL21 (DE3) cells were transformed with pLE601 with or (as a control) without the csCyDisCo plasmid (pLE577) (25). Cells were plated on lysogeny broth (LB) agar supplemented with kanamycin (50 µg/mL) with (when co-transforming with pLE577) or without chloramphenicol (30 µg/mL). A single colony was picked to inoculate the LB medium containing the same type and concentration of antibiotic as used on the LB agar plates. The overnight culture was incubated for *∼*16 hours at 37°C at 200 rpm in an orbital shaker.

For initial small-scale expression tests (50 mL), LB medium containing appropriate antibiotics and supplemented with 0.05% glucose was inoculated with 2% overnight culture and grown at 37°C with shaking at 200 rpm until the desired OD_600_ of 0.6-0.8 was reached. Expression was induced by adding isopropyl ß-D-1-thiogalactopyranoside (IPTG) to a final concentration of 1 mM, and the cultures grown for 18 hours at 25°C with shaking at 200 rpm to allow protein expression. Large-scale expression (1 L culture volume) was performed in auto-induction media prepared as described previously (67). Briefly, terrific broth medium containing kanamycin (100 µg/mL) and/or chloramphenicol (30 µg/mL) was supplemented with sterilized stocks of the following: 0.05% glucose, 0.2% lactose, 50 mM KH_2_PO_4_/Na_2_HPO_4_, 50 mM NH_4_Cl, 50 mM Na_2_SO_4_, 0.1 mM FeCl_3_, 2 mM MgSO_4_, 0.1 mM CaCl_2_ and 1 x metal mix (203 g/L MgCl_2_ 6·H_2_O, 2.1 g/L CaCl_2_ 2·H_2_O, 2.7 g/L FeSO_4_ 7·H_2_O, 20 mg/L AlCl_3_ 6·H_2_O, 10 mg/L CoSO_4_ 7·H_2_O, 2 mg/L KCr(SO_4_)_2_ 12·H_2_O, 2 mg/L CuCl_2_ 2·H_2_O, 1 mg/L H_3_BO_4_, 20 mg/L KI, 20 mg/L MnSO_4_ H_2_O, 1 mg/L NiSO_4_ 6·H_2_O, 4 mg/L Na_2_ MoO_4_ 2·H_2_O, 4 mg/L ZnSO_4_ 7·H_2_O, 21 g/L citric acid monohydrate). Cultures were grown in 37°C at 200 rpm until OD_600_ reached 0.8, at which point cells were moved to 25°C for expression performed with shaking at 200 rpm for 18 hours.

### Harvest and clarification of bacterial cultures

Induced cultures were harvested by centrifugation at 5,000 g for 20 min. The cell pellets were resuspended in 5 mL lysis buffer (50 mM Tris, pH 8, 300 mM NaCl, 20 mM imidazole) per gram pellet. Cell resuspensions were supplemented with ∼12 units Benzonase Nuclease (Merck Millipore)/L culture to minimize viscosity due to the presence of nucleic acids post-lysis. Cell lysis was performed using a UP200S ultrasonic processor (Hielscher) keeping the cells on ice throughout. Cells were lysed with 8 x 30-sec pulses at 90% power with 30 sec rests between each pulse. Cell debris was pelleted by centrifugation at 30,000 g for 45 minutes. The cleared lysates were filtered through 0.45 µm syringe filters and transferred to fresh tubes, while the pellets were resuspended in an equal volume lysis buffer containing 8 M urea for SDS-PAGE analysis.

### Protein purification

Ub-His_10_-Mu8.1 was affinity purified from the clarified lysate on an ÄKTA START system equipped with a 5 mL prepacked HisTrap HP (Cytiva) column equilibrated in lysis buffer. The lysate was applied to the column and washed with approximately 20 column volumes (CV) of lysis buffer before elution of Ub-His_10_-Mu8.1 with a gradient of 0 to 100% elution buffer (50 mM Tris, pH 8, 300 mM NaCl, 400 mM imidazole) applied over 20 CVs. Pooled fractions were dialyzed twice against 2 L anion exchange (AEX) buffer (50 mM NaH_2_PO_4_/Na_2_HPO_4_, pH 6.8, 20 mM NaCl). Anion exchange chromatography was performed on an ÄKTA Pure system equipped with a 10/100 Tricorn column (Cytiva) packed with Source 15Q ion exchange resin (Amersham Biosciences, GE Healthcare) equilibrated in AEX buffer. Ub-His_10_-Mu8.1 was eluted using a gradient from 15% to 50% AEX elution buffer (50 mM NaH_2_PO_4_/Na_2_HPO_4_, pH 6.8, 1 M NaCl) developed over 6 CVs.

The Ub-His_10_-Mu8.1 fusion protein was cleaved using His_6_-tagged TEV protease, expressed, and purified as described previously (25). A molar ratio Ub-His_10_-Mu8.1:His_6_-TEV protease of 1:20 was used. To avoid reducing the disulfides in Mu8.1, His_6_-TEV protease – pre-activated with 2 mM dithiothreitol (DTT) for 30 min at room temperature – was diluted to ∼0.002 mM DTT by three rounds of dilution/concentration in an Amicon Ultra 15 mL 3K Centrifugal Filter (Merck Millipore). TEV protease cleavage was performed overnight at room temperature.

To remove uncleaved Ub-His_10_-Mu8.1, the Ub-His_10_ tag, and His_6_-TEV, the cleavage mixture was applied to a gravity flow column packed with 8 mL TALON cobalt resin (Takara) equilibrated in AEX buffer. The flow-through and the first wash fraction were collected. The presence of cleaved Mu8.1 in each fraction was investigated by analysis on 15% tricine SDS-PAGE gels, and the protein-containing fractions pooled. The cleaved Mu8.1 was subjected to size exclusion chromatography on a Superdex 75 Increase 10/300 GL column (Cytiva) equilibrated in 200 mM NH_4_HCO_3_ buffer, pH 7.8. Fractions containing purified Mu8.1 were pooled and lyophilized.

### Analytical gel filtration

To analyze the oligomeric state of Mu8.1, the protein was analyzed at two concentrations, 100 µm and 1 µm, on a Superdex 75 Increase 10/300 GL column (Cytiva) equilibrated in 10 mM NaPi, pH 7.8, 150 mM NaCl at a flowrate of 0.5 mL/min. An excess of the protein was loaded onto a 100 µL loop to ensure analysis of the same volume in each run.

### SDS-PAGE analysis

Samples from bacterial expression and subsequent purification steps were separated on 15% glycine SDS-PAGE or 15% tricine SDS-PAGE gels (68). Where indicated, reduced samples were treated with 40 mM DTT. Protein bands were visualized with Coomassie Brilliant Blue and images were recorded with a BioRad Chemidoc Imaging System. For western blotting, proteins separated by SDS-PAGE were transferred to a PVDF membrane (Immobilon-P, Merck Millipore) in a Mini Trans-Blot (Bio-Rad) transfer system. A mouse monoclonal His-tetra (Qiagen) antibody (1:1000 dilution) was used in combination with a horse-radish peroxidase-conjugated *α*-mouse secondary antibody (Pierce) (1:100,000 dilution). Chemiluminescence detection was performed using ECL Select Peroxide and Luminol solutions (GE Healthcare) according to the manufacturer’s directions.

### Concentration determination

Concentrations were determined by measuring absorbance at 280 nm and using the theoretical extinction coefficient provided by the Expasy ProtParam tool available through the Expasy bioinformatics resource web portal (69). Concentrations used for bioassays, constellation pharmacology, and electrophysiology assumed a monomeric state of Mu8.1.

### Determination of molecular mass in solution by small angle x-ray scattering (SAXS)

Prior to SAXS analysis, Mu8.1 was run through a Superdex 75 Increase 10/300 GL column to ensure a monodisperse sample. The protein was subsequently dialyzed into a solution containing 150 mM NaCl and 10 mM NaPi, pH 7.8, and the dialysis buffer was reserved for measurement of background scattering. Immediately preceding SAXS measurements any precipitates were removed from the sample by centrifugation at 20,000 g for 15 min at 4°C. Six dilutions were prepared ranging from 0.5 mg/mL to 12.4 mg/mL (49 µM to 1.2 mM) in dilution buffer. SAXS data was collected by the beamline staff at CPHSAXS using a Xenocs BioXolver L equipped with a liquid gallium X-ray source (λ=1.34 Å). A sample-to-detector distance of 632.5 mm was used, corresponding to a Q-range of 0.013 – 0.5 Å^-1^. Samples and buffer were measured at room temperature, and automatic loading was performed robotically from a 96-well plate. Data were collected as a minimum of 10 frames with 120 sec exposure per frame. The longer exposure time was to account for the lower concentration and the expected presence of multimeric species. Data reduction and primary analysis were performed using RAW (70), SVD, and oligomer analysis performed with OLIGOMER (ATSAS program package) (71) and scattering curves plotted in Matlab R2020b.

### X-ray crystallography

Freeze-dried Mu8.1 was dissolved in Milli-Q water to a concentration of 5 mg/mL. Crystallization screening experiments were performed with the Structure screen II (Molecular Dimensions, MD1-02) and the Index screen (Hampton Research, HR2-144) by the hanging drop vapor diffusion method. The crystal drops were mixed using 1 µL of protein and 1 µL precipitant solution as hanging drops on siliconized glass cover-slides and equilibrated against 500 µL of precipitant solution in a 24-well plate setup. Wells were sealed with immersion oil (Sigma-Aldrich) and incubated at 21°C. Initial crystals of Mu8.1 appeared in several conditions in a few days to weeks. Diffracting crystals were obtained from the Structure screen II condition 38 (0.1 M NaOAc pH 4.6, 0.1 M CdCl hemi(pentahydrate), 30% v/v PEG400), abbreviated as Mu8.1_38, and from the Index screen condition 59 (0.1 M HEPES pH 7.5, 0.02 M magnesium chloride hexahydrate, 22% w/v polyacrylic acid sodium salt 5,100), abbreviated as Mu8.1_59.

Crystals were harvested using mounted CryoLoops (Hampton Research) and flash-cooled in liquid nitrogen. Cryoprotection was performed by quickly dipping the crystal in ∼17 % ethylene glycol (EG) prepared by mixing 1 µL 50% EG with 2 µL of the reservoir condition specific to each crystal condition. Both native crystals and iodide-soaked crystals were prepared from all three conditions. Single iodide crystals (Sigma-Aldrich) were added to the above-mentioned cryo condition for each of the crystal conditions. Crystals from each condition were transferred to the iodide-containing cryo conditions and left to soak 5-10 sec before harvesting them.

### Data collection

Flash-cooled crystals were shipped to the beamline for remote data collection. Data were collected at 100K on a PILATUS detector at BioMax (MAX-IV, Lund, Sweden). A full sweep of 360° data was collected with an oscillation degree of 0.1°, with 0.050 s exposure at 12700 eV and 7000 eV. Complete data set was processed from 360° (3600 images) with the x-ray beam reduced to 5% intensity.

### Data processing

Native data was collected for Mu8.1_59 at 12700 eV and only Mu8.1_38 showed an anomalous signal from the data collected at 7000 eV. All data was processed with xia2 using the 3dii pipeline (72, 73) (Table 1).

**Table 1.** Data collection and refinement statistics.

| Name (PDB ID) | Mu8.1_59* (7PX2) | Mu8.1_38* (7PX1) |
| --- | --- | --- |
| Wavelength (Å) | 1.771 | 1.771 |
| Resolution range (Å) | 48.97 - 2.12 (2.196 - 2.12) | 36.62 - 2.33 (2.413 - 2.33) |
| Space group | P 1 21 1 | P 21 21 2 |
| a, b, c (Å) | 68.8, 88.4, 69.3 | 49.3, 109.3, 34.3 |
| $\alpha$, $\beta$, $\gamma$ (°) | 90, 116.71, 90 | 90, 90, 90 |
| Total reflections | 258079 (14423) | 103396 (10360) |
| Unique reflections | 41056 (3498) | 8423 (811) |
| Multiplicity | 6.3 (4.1) | 12.3 (12.8) |
| Completeness (%) | 97.3 (83.2) | 99.7 (99.9) |
| Mean I/sigma (I) | 9.75 (1.21) | 7.62 (1.08) |
| Wilson B-factor(Å<sup>2</sup>) | 43.38 | 47.60 |
| R-merge | 0.091 (0.81) | 0.21 (>1) |
| R-meas | 0.099 (0.92) | 0.21 (>1) |
| R-pim | 0.039 (0.44) | 0.064 (0.38) |
| CC1/2 | 0.99 (0.68) | 0.99 (0.60) |
| CC* | 0.99 (0.89) | 0.99 (0.87) |
| Reflections used in refinement | 41043 (3498) | 8408 (812) |
| Reflections used for R-free | 2082 (167) | 446 (42) |
| **R-work** | 0.21 (0.38) | 0.26 (0.37) |
| **R-free** | 0.23 (0.38) | 0.30(0.48) |
| **CC (work)** | 0.95 (0.80) | 0.92 (0.81) |
| **CC (free)** | 0.96 (0.80) | 0.87 (0.57) |
| **Number of non-hydrogen atoms** | 4299 | 1386 |
| **Macromolecules** | 4086 | 1346 |
| **Ligands** | 192 | 13 |
| **Solvent** | 123 | 27 |
| **Protein residues** | 510 | 168 |
| **r.m.s.d (bonds) (Å)** | 0.006 | 0.003 |
| **r.m.s.d (angles) (°)** | 1.03 | 0.49 |
| **Ramachandran favored (%)** | 99.00 | 97.56 |
| **Ramachandran allowed (%)** | 1.00 | 1.83 |
| **Ramachandran outliers (%)** | 0.00 | 0.61 |
| **Rotamer outliers (%)** | 0.64 | 0.65 |
| **Clash score** | 5.81 | 5.72 |
| **Average B-factor (Å<sup>2</sup>)** | 58.2 | 53.6 |
| **Macromolecules** | 58.00 | 53.40 |
| **Ligands** | 64.69 | 78.83 |
| **Solvent** | 60.20 | 54.09 |
| **Number of TLS groups** | 18 | 10 |

The phases for Mu8.1_38 were experimentally determined using autosol in the PHENIX package (74) with 13 iodide sites identified and an initial Figure of Merit (FOM) of 0.4. Density resembling helical structures were visible in the electron density map. The following AutoBuild wizard within the PHENIX package (74) was able to build a preliminary structure with the main helices in place. This structure was used as initial search model for molecular replacement and performed with the program Phaser (75) against the highest resolution dataset Mu8.1_59.

All structures were manually refined using phenix.refine (74), and final model building was performed in Coot (76). Data collection and refinement statistics are summarized in Table 1. Molecular graphics were presented with the PyMOL Molecular Graphics System, Version 2.2r7pre, Schrödinger, LLC. Electrostatic potentials were modelled using the Adaptive Poisson-Boltzmann Solver (APBS) plugin in PyMOL2.3 (77).

### Structure search and topological comparisons

Structures similar to Mu8.1 were identified using PDBeFold at the European Bioinformatics Institute (https://www.ebi.ac.uk/msd-srv/ssm/) (78, 79). Monomeric Mu8.1 was used as query molecule to identify similar structures with a minimum acceptable match set to 60% or higher from the entire PDB database. Structural overlays were generated with the CLICK structural alignment tool (http://cospi.iiserpune.ac.in/click/) selecting ‘CA’ as representative atoms and visualized using the PyMOL Molecular Graphics System (80), or executed manually in PyMol.

### Zebrafish bioassay

Adult wild-type zebrafish, *Danio rerio,* were raised under standard conditions with a 14/10 h light/darkness cycle. Intramuscular injections were approved by the University of Utah Institutional Animal Care and Use Committee (IACUC). The test (n=3) and control (n=3) fish (originating from separate clutches) were subjected to the exact same procedures. Prior to the injection, adult zebrafish (3-6 months old) were placed in a rectangular open arena and allowed to swim for 20 min to acclimate to the environment. After the conditioning period, 5 nmols of lyophilized Mu8.1 reconstituted in 12 µL of normal saline solution (NSS) or 12 µL of NSS were injected at the caudal end of the fish below the dorsal fin using a 31-gauge syringe. The animals were quickly transferred to the same rectangular arena and allowed to swim for 4 hours after the injection. The behavior of the animals was recorded at 20 frames per second under controlled light conditions for 20 min intervals.

### Video analysis of zebrafish behavior

All videos were analyzed with two custom-built python scripts. First, the videos were transformed into binary files, and background subtraction was performed (source code: https://github.com/pflorez/Ruby-Tracker/blob/master/bsub.py). The binarized output was used to track individual free-swimming fish (source code: https://github.com/pflorez/Ruby-Tracker/blob/master/ruby_tracker.py).

### Mouse bioassay

Mouse intracranial injections were approved by the University of Utah Institutional Animal Care and Use Committee (IACUC). 10 nmols of lyophilized Mu8.1 were reconstituted in 20 µL of normal saline solution (NSS; 0.9% NaCl) and injected intracranially into 12-day old Swiss Webster mice (Simonsen Laboratories; n=3). Negative controls (n=2) were injected with 20 µL of NSS. Following injection, the mice were placed in individual observation arenas where their behavior was observed for 4 hours. All animals were kept overnight.

### Constellation pharmacology

Primary cell cultures were dissociated from calcitonin gene-related peptide (CGRP)-GFP mice, STOCK Tg(Calca-EGFP)FG104Gsat/Mmucd, ages 34-38 days old as described previously (11). In brief, lumbar dorsal root ganglia from vertebrae L1-L6 were dissected, trimmed and treated with 0.25% trypsin for 20 min. Following trypsinization, the DRGs were mechanically triturated using fired polished pipettes and plated in poly-l-lysine-coated plates. All plated cells were kept overnight at 37°C in a minimal essential medium supplemented with 10% fetal bovine serum, 1X penicillin/streptomycin, 10 mM HEPES, and 0.4% (w/v) glucose. One hour before the experiment, the dissociated cells were loaded with 4 µM Fura-2-AM dye (Sigma-Aldrich) and kept at 37°C. During each experiment, all dissociated cells were exposed to different pharmacological agents utilizing an automatic perfusion system and were imaged at 340/380 nm at 2 frames per second. In brief, cells were incubated with the pharmacological agents for 15 seconds followed by 6 consecutive washes and a 5-minute incubation period with extracellular solution for controls or Mu8.1. Five different pharmacological agents were used for cell classification: mustard oil (allyl isothiocyanate, AITC) at 100 µM, menthol at 400 µM, capsaicin at 300 nM, K^+^ at 25 and 40 mM, and conotoxin kM-RIIIJ at 1 µM as described previously (11). SNX-482 (Alomone Labs) was used at 100 nM. At the end of each experiment, all cells were incubated for 7 minutes with a 2.5 µg/mL Alexa-Fluor 647 Azolectin B4 (IB4). All data acquisition was performed using the Nikon NIS-Elements platform. Cellprofiler (81) was used for region of interest (ROI) selection and a custom-built script in python and R was used for further data analysis and visualization^1^.

### Statistical analysis of cellular calcium imaging

We utilized a Min-Max normalization (f(x)) to assess the effects of Mu8.1 on the peak height of the calcium signal induced by high concentrations of potassium using the formula:

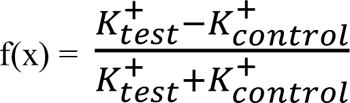

where K^+^ test is the peak height after the incubation with Mu8.1 and K^+^ control is the peak height before the incubation with Mu8.1 (11). If f(x) = 0, this suggests that the conotoxin did not affect the Ca^2+^ concentration in the cytosol induced by the high concertation of potassium. An f(x) > 0 indicates that the conotoxin increased the cytosolic calcium concentration after a high potassium concentration, resulting in an amplification. Finally, if f(x)<0, the conotoxin decreased calcium concentration in the cytosol resulting in a calcium block. After calculating the f(x) for all sensory neuron cell types, we performed a two-tailed t-test to assess if the f(x) values calculated for every cell type were significantly different from 0.

### Electrophysiology

Automated patch clamp (APC) recordings were performed in a Patchliner Octo (Nanion Technologies GmbH, Munich, Germany) equipped with two EPC-10 quadro patch-clamp amplifiers (HEKA Electronics). PatchControlHT (Nanion) was used for cell capture, seal formation and establishment of the whole-cell configuration, whilst voltage was controlled, and currents sampled with PatchMaster (HEKA Electronik). Recordings were performed under the whole-cell configuration using single-hole planar NPC-16 chips (resistance of ∼2.5 MΩ) at room temperature. Stably transfected cell lines (D.J. Adams collection, IHMRI-UOW) were cultured according to manufacturer’s instructions and detached using TrypLE. Cells were resuspended in cold external recording solution and kept in suspension by automatic pipetting at 4°C. The extracellular solution used for Kv1, hERG and Nav1 recordings contained (in mM): 140 NaCl, 5 KCl, 2 CaCl_2_, 2 MgCl_2_, 10 glucose, and 10 HEPES (pH 7.4 with NaOH, 298 mOsmol/kg). Kv1 and hERG intracellular solution (in mM): 60 KF, 70 KCl, 10 EGTA, 10 glucose, and 10 HEPES (pH 7.2 with KOH, 285 mOsmol/kg). Peak I_K_ currents for Kv1.1-3: 500 ms test pulse to 20 mV (Vh = *−*120 mV; 0.1 Hz). hERG I_K_: 1 s prepulse to +40 mV, was followed by 200 ms test pulse to −40 mV (Vh = *−*80 mV; 0.1 Hz). Nav1 intracellular solution (mM): 60 CsF, 60 CsCl, 10 NaCl, 10 EGTA, and 10 HEPES (pH 7.2 with CsOH, 285 mOsmol/kg). Nav currents were elicited by 10 ms test pulses to *−*10 mV with a 1 sec prepulse to *−*120 mV (Vh = *−*90 mV; 0.1 Hz). APC of Cav2: extracellular solution (in mM): 135 NaCl, 4 KCl, 10 BaCl_2_, 1 MgCl_2_, 5 glucose, and 10 HEPES (pH 7.4 with NaOH, 298 mOsmol/kg) and intracellular solution (in mM): 90 CsSO_4_, 10 CsCl, 10 NaCl, 10 EGTA, and 10 HEPES (pH 7.2 with CsOH, 285 mOsmol/kg). Peak calcium currents were measured upon 50 ms step depolarization to 10 mV (Vh = *−*80 mV; 0.1 Hz). Recordings where seal resistance (SR) was >500 MΩ and access resistance was <3xSR were considered acceptable. Chip and whole-cell capacitance were fully compensated, and series resistance compensation (70%) was applied via Auto Rs Comp function. Recordings were acquired with PatchMaster (HEKA Elektronik, Lambrecht/Pfalz, Germany) and stored on a computer running PatchControlHT software (Nanion Technologies GmbH, Munich, Germany).

Manual patch-clamp (MPC) was performed on human embryonic kidney (HEK293T) cells containing the SV40 Large T-antigen cultured and transiently transfected by calcium phosphate method as reported previously (82). In brief, cells were cultured at 37°C, 5% CO_2_ in Dulbecco’s Modified Eagle’s Medium (DMEM, Invitrogen Life Technologies, VIC, Australia), supplemented with 10% fetal bovine serum (FBS, Bovigen, VIC, Australia), 1% GlutaMAX and penicillin-streptomycin (Invitrogen). The human orthologues of Cav3.1, Cav3.2 ad Cav3.3 channels were co-transfected with GFP for identification of positive transfectants. cDNAs encoding hCav3.1 (kindly provided by G. Zamponi, University of Calgary), hCav3.2 (a1Ha-pcDNA3, Addgene #45809) and hCav3.3 (a1Ic-HE3-pcDNA3, Addgene #45810) were a kind gift from E. Perez-Reyes (University of Virginia). MPC experiments employed a MultiClamp 700B amplifier, digitalized with a DigiData1440 and controlled using Clampex11.1 software (Molecular Devices, CA, USA). Recordings of I_Ca_ through hCav3.1-3 were performed using an extracellular solution containing (in mM): 110 NaCl, 10 CaCl_2_, 1 MgCl_2_, 5 CsCl, 30 TEA-Cl, 10 D-Glucose and 10 HEPES (pH 7.35 with TEA-OH, 305 mOsmol/kg). Pipettes were pulled from borosilicate glass capillaries (GC150F-15, Harvard Apparatus, MA, USA), fire polished to a final resistance of 1-3 MΩ and filled with intracellular solution (in mM): 140 KGluconate, 5 NaCl, 2 MgCl_2_, 5 EGTA and 10 HEPES (pH 7.2 with KOH, 295 mOsmol/kg). Peak currents were measured upon stimulation using 50 ms test pulses to −20 mV from a holding potential (Vh) of −90 mV and pulsed at 0.2 Hz. Whole-cell currents were sampled at 100 kHz and filtered to 10 kHz, with leak and capacitive currents subtracted using a P/4 protocol, and 60-80% series resistance compensation.

### Data analysis of electrophysiology experiments

APC analysis was performed using Igor Pro-6.37 (WaveMetrics Inc.). Cav2.3 peak currents measured in the presence of increasing Mu8.1 concentrations (I_Mu8.1_) were divided by the current in control conditions (I_Ctr_) to generate a concentration response curve that was fit with a Hill equation of the form:

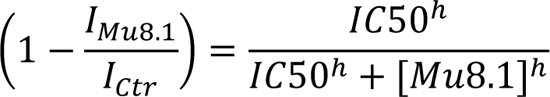

Where IC_50_ is the half-maximal inhibitory concentration, and h is the Hill coefficient (nH).

For ease of comparison, IC_50_ values were calculated from fractional block for the other voltage gated channels that were screened at a single Mu8.1 concentrations according to following equation:

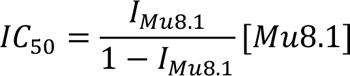

## Acknowledgements

We acknowledge the MAX IV Laboratory for time on Beamline Biomax under Proposal 20190334, and the University of Copenhagen, Small Angle X-ray facility, CPHSAXS (https://drug.ku.dk/core-facilities/cphsaxs/), funded by the Novo Nordisk Foundation (grant no. NNF19OC0055857). We thank Dr. Uwe Müller for assistance during the data collection, and Cecilie L. Søltoft for expert technical assistance. Radioligand binding and GPCR binding assays were generously provided by the National Institute of Mental Health’s Psychoactive Drug Screening Program, Contract # HHSN-271-2018-00023-C (NIMH PDSP). The NIMH PDSP is Directed by Dr. Bryan L. Roth at the University of North Carolina at Chapel Hill and Project Officer Jamie Driscoll at NIMH, Bethesda MD, USA.

## Author contributions

Conceptualization, C.M.H., H.S.-H. and L.E.; Data curation, P.F.-S. and R.K.F.-U.; Formal Analysis, P.F.-S., E.M., T.L.K., P.S.T., J.P.M. and R.K.F.-U.; Funding acquisition, B.O., H.S.-H., J.P.M., J.R.M., D.J.A., A.S.K. and L.E.; Investigation, C.M.H., L.D.K., L.G.D., M.W., E.M., T.L.K., P.S.T., J.P.M., J.R.M., H.S.-H., P.F.-S. and R.K.F.-U.; Project administration, H.S.-H., C.M.H. and L.E.; Supervision, B.O., J.P.M., H.S.-H., A.S.K. and L.E.; Visualization, C.M.H., L.G.D., P.F.-S. and R.K.F.-U.; writing—original draft preparation, C.M.H., B.O., L.G.D., E.M., T.L.K., H.S.-H., J.P.M., P.F.-S., R.K.F.-U. and L.E.; writing—review and editing, C.M.H., B.O., E.M., H.S.-H., J.P.M., J.R.M., D.J.A., P.F.-S., R.K.F.-U. and L.E. All authors have read and agreed to the published version of the manuscript.

## Database depositions

Nucleotide sequence data for Mu8.1 and Mu8.1ii have been submitted to GenBank with accession numbers ON755370 and ON755371, respectively. The atomic coordinates the structure-factor amplitudes for Mu8.1_38 and Mu8.1_59 have been deposited with the Protein Data Bank under accession numbers 7PX1 and 7PX2, respectively.

## Footnotes

^1^The package used to visualize, analyze, and deploy models for constellation pharmacology experiments is available at: https://github.com/leeleavitt/procPharm.

## Abbreviations

AEX: anion exchange

AITC: allyl isothiocyanate derived from mustard oil

CNGB: China National Genebank

APC: automated patch clamp

CD: circular dichroism spectroscopy

CGRP: calcitonin gene-related peptide

C-LTMRs: C-low threshold mechanoreceptors

CNS: central nervous system

csPDI: conotoxin-specific PDI

CV: column volume

DDBJ: DNA Databank of Japan

DMEM: Dulbecco’s Modified Eagle’s Medium

DRG: dorsal root ganglia

DTT: dithiothreitol

ER: endoplasmic reticulum

FDA: U.S. Food and Drug Administration

FOM: figure of merit

GFP: green fluorescent protein

GPCR: G protein-coupled receptor

HEK293T: human embryonic kidney cells

hPDI: human PDI

IACUC: Institutional Animal Care and Use Committee

IB4: Alexa-Fluor 647 Azolectin B4

IMAC: immobilized metal affinity chromatography

IPTG: isopropyl ß-D-1-thiogalactopyranoside

LB: lysogeny broth

MALDI - TOF: matrix-assisted laser desorption-ionization-time of flight

MPC: manual patch clamp

NCBI: National Center for Biotechnology Information

NSS: normal saline solution

PDI: protein-disulfide isomerase

RMSD: root mean square deviation

RP-HPLC: reversed-phase high pressure liquid chromatography

SAPLIP: Saposin like proteins

SAXS: small angle x-ray scattering SVD

SDS-PAGE: sodium dodecyl sulfate-polyacrylamide gel electrophoresis

SLC: saposin-like conotoxin

SR: seal resistant

TEV: tobacco etch virus

Ub: ubiquitin

Ub-His_10_: Ub containing 10 consecutive histidines.

