## Supplementary Information for "Identification of a sensory neuron Cav2.3 inhibitor within a new superfamily of macro-conotoxins"

### Materials and Methods

**Transcriptome mining.** The transcriptomes of 44 species of cone snails publicly available from NCBI, DDBJ, and CNGB repositories were mined with Mu8.1 and coni-ikot-ikot sequences as queries (**Table S1**). The transcriptomes were assembled as previously described (1). Using tBLASTn with e<1e-3 we identified 496 transcripts with sequence similarity to Mu8.1 and con-ikot-ikots (Supplementary file A).

**CLANS clustering analysis.** To investigate the relationship of the identified toxin sequences, we performed a CLANS clustering analysis (2) using the web tool https://toolkit.tuebingen.mpg.de/tools/clans. CLANS performs an all-against-all BLASTp and retrieves the negative logarithm of the resulting p-values < 1e-6, which are used as attractive forces between the nodes representing the sequences. The additional uniformly repulsive field between all nodes forces the nodes with the lowest blast p-values to cluster together. The BLAST p-values were calculated using the BLOSUM62 matrix. The clustering was first done in 3D for 20,000 rounds and subsequently in 2D for an additional 10,000 rounds, at which point the nodes converged to an equilibrium. The resulting clans-file is supplied in Supplementary file B and can be viewed using the command: java -Xmx4G -jar clans.jar -load Supplementary_file_B.clans.jar

**UTR sequence alignment.** We aligned the complete transcripts of Mu8.1, an additional SLC transcript, and two random transcripts from each of the other three toxin clusters using MAFFT L-Ins-I v7.487 and colored the alignment using https://www.bioinformatics.org/sms2/color_align_cons.html. Fig. S3 shows the 5' UTR and initial segment of the ORF.

**Gene structure and intron sequence alignment.** We identified two transcripts from *Conus ventricosus* and six from *Conus betulinus* that could be mapped in their entire length to the respective genomes. We extracted the 100 intronic nucleotides flanking the ORF encoding exons (Supplementary file C) and aligned the 5' and 3' regions separately using MAFFT L-Ins-I v7.487. Supplementary file C shows the percentage sequence identity of the intron regions. The transcripts were mapped onto the genome using Splign v5 (3) to identify the location and phase of the introns.

**Determination of protein oxidation state by MALDI-TOF mass spectrometry (MS)**. Mu8.1 (500 µL) was first desalted by reversed-phase high-performance liquid chromatography (RP-HPLC) chromatography. Trifluoroacetic acid (TFA) was added to a final concentration of 0.1%. The acidified sample was centrifuged at 16,100 g for 15 min and RP-HPLC was performed on an ÄKTA purifier 900 system equipped with a Kromasil C18 column (4.6 mm x 150 mm, 5 µm) using solvent A (5% ethanol, 0.1% TFA) and solvent B (90% ethanol, 0.1% TFA). Elution was executed as a 0% to 75% gradient over 35 mL at a flowrate of 1 mL/min. Protein-containing fractions were analyzed by MALDI-TOF MS using an Autoflex Smartbeam III instrument (Bruker) calibrated by external calibration (Peptide calibration standard I; Bruker Daltronics). Samples for analysis were mixed with alpha-cyano-4-hydroxycinnamic acid matrix prepared with 70% acetonitrile, 0.1% TFA, spotted on a stainless-steel target plate and analyzed in positive reflectron mode.

**Circular Dichroism (CD) spectroscopy**. CD spectroscopy was performed on a JASCO J-810 CD Spectropolarimeter. Mu8.1 was dissolved in 10 mM sodium phosphate buffer, pH 8 to a final concentration of 10 µm Mu8.1 in 200 µL buffer. CD spectra were recorded in a 0.1 cm cuvette at 25°C in the wavelength range of 190 nm to 260 nm using a data pitch of 0.1 nm and a bandwidth of 1.0 nm. The final spectra were obtained by averaging 10 spectra recorded at a scan rate of 10 nm/min and the baseline (buffer only) was subtracted. The measured ellipticity was then converted to residual molar ellipticity and the data were visualized using Matlab (MathWorks). Deconvolution of the CD spectrum was calculated using BestSel, a web-based deconvolution tool (4).

**Intracellular $Ca^{2+}$ imaging for determination of the effect of Mu8.1 on ionotropic glutamate receptor subtype AMPA receptors**. Human embryonic kidney (HEK) 293T cells (American Type Culture Collection, Manassas, VA) were transiently transfected with rGluA2(Q)i plasmid DNA using LipdoD293™ DNA In Vitro Transfection reagent by following the protocol supplied by the manufacturer. Briefly, HEK293T cells in suspension were mixed with DNA/transfection complex (formed by mixing plasmid DNA, LipdoD293 reagent, and DMEM in a 1:2:25 ratio) and plated into poly-D-lysine–

coated Falcon black clear-bottom 96-well plates (Corning, Corning, NY) to a final density of approximately 20,000 cells and 0.025 µg plasmid DNA per well. The competitive antagonist CNQX was added at a final concentration of 20 µM to protect against glutamate-induced cytotoxicity. Cells were incubated for 2 days after transfection before experiments. Intracellular calcium concentration, serving as an indirect measure of receptor activation, upon agonist application, were measured as a change in fluorescence of Fluo-8 AM using a FlexStation I plate reader (Molecular Devices). The experiment was performed in the presence and absence of cyclothiazide (CTZ), a positive allosteric modulator of AMPA receptors, known to block receptor desensitization and thus increase AMPA receptor current (5). On the day of experiments, transfected cells were washed three times in Phosphate Buffered Saline with $Ca^{2+}$ and $Mg^{2+}$ (in mM: 137 NaCl, 2.7 KCl, 10 $Na_2HPO_4$, 2 $KH_2PO_4$, 0.1 $CaCl_2$, 0.5 $MgCl_2$, pH 7.4) and loaded with a solution containing 2 µM Fluo-8 AM fluorescent indicator dye (dissolved in plain DMEM) and incubated for 30 min at 37 °C. Excess loading dye was removed by washing three times in FLUO buffer containing (in mM) 140 choline chloride, 5 KCl, 1 $MgCl_2$, 10 $CaCl_2$, 10 HEPES (pH 7.4). Cells were pre-incubated with 50 µl FLUO buffer containing various concentrations of Mu8.1 in the presence and absence of CTZ for 30 min at room temperature. Changes in dye fluorescence upon addition of a saturating agonist solution with various concentrations of Mu8.1 in the presence and absence of CTZ were measured at 538 nm using excitation at 485 nm. Final concentrations used in the experiment were 1 mM glutamate (agonist), 100 µM CTZ and 0.05, 0.5, 5 µM Mu8.1. The experiment was performed in quadruplicate wells.

**Initial receptor and GPCRome screening**. Preliminary functional screenings were performed through the National Insititute of Mental Health's Psychoactive Drug Screening Program (PDSP), contract #HHSN-271-2018-00023-C (NIMH PDSP). The NIMH PDSP is directed by Bryan L. Roth at the University of North Carolina at Chapel Hill and Project Officer Jamie Driscoll at NIMH, Bethesda, MD, USA. Detailed protocols for all assays can be found in the NIMH PDSP Assay Protocol Book accessed via the PDSP website http://pdsp.med.unc.edu/.

**Radioligand binding assays**. A primary binding assay was performed on membrane preparations derived from cell lines transiently or stably expressing 52 different receptors (6). Evidence for interaction was based on the inhibition of a reference ligand-binding signal. Secondary binding assays were performed only when Mu8.1 had a signal inhibition greater than 50%. For the primary binding assays, Mu8.1 was applied at a single concentration (10 μM) in quadruplicate in 96-well plates. In secondary binding assays, Mu8.1 was tested in triplicate at eleven concentrations (0.1, 0.3, 1, 3, 10, 30, 100, 300 nM, 1, 3, 10 μM). Both primary and secondary binding assays were carried out in a final volume of 125 μL per well. The "hot" ligand was usually applied at a concentration close to the Kd. Total binding and nonspecific binding were determined in the absence and presence of 10 μM of the appropriate reference compound, respectively. Reactions were stopped by vacuum filtration onto 0.3% polyethyleneimine (PEI) soaked 96-well filter mats using a 96-well Filtermate harvester, followed by three washes with cold wash buffers. Scintillation cocktail was then melted onto the microwave-dried filters on a hot plate and radioactivity was counted in a Microbeta counter.

**GPCR binding assays**. Mu8.1 was also tested on a panel of GPCRs using the PRESTO-Tango system (7). This assay employs HTLA cells, a HEK293T cell-derived stable cell line expressing a human β-arrestin2-TEV protease fusion and a tetracycline-controlled transactivator (tTA)-dependent firefly luciferase reporter gene. Briefly, cells were plated in 384-well plates and incubated overnight. The cells were then transfected with the receptor constructs (318 target receptors) and incubated overnight at 37°C. Following overnight transfection, cells were treated with 10 μM Mu8.1 and incubated overnight at 37°C. Mu8.1 and the media were then removed and Bright-Glo (Promega) reagent added to determine luciferase activity. Results are presented as fold of the average basal activity. Activity between 0.5 to 2.0-fold of basal is considered normal and not warranting further testing. 100 nM quinpirole stimulation of dopamine receptor DRD2 was used as an assay control.

**Analysis of saposin domain-containing proteins.** 13,594 saposin domain-containing proteins were downloaded from Uniprot with the search terms "annotation:(type:"positional domain" saposin)" and clustered with cd-hit v4.8.1 using -

c 0.8. The reduced dataset was used as queries against the *C. ventricosus* transcriptome and genome using tBLASTn with an e-value below 1e-10. This search identified five *C. ventricosus* proteins with saposin-like domains (Supplementary file D). The gene structures of the *C. ventricosus* saposin domain-containing proteins were determined in a similar way as for the toxins.

**AlphaFold structure prediction.** Structural prediction of a randomly selected toxin from Cluster 1 (Supplementary file A (>14718X8.C.litteratus.TRINITY_DN2335_c0_g2_i1_Entry:5131.conotoxin.Conikotikot_len:998_tpm:5563.29 45) and (8)) was generated using the AlphaFold2 (9) implementation in the ColabFold notebook running on Google Colaboratory (10). The predicted structure was produced using the default settings and a custom multiple sequence alignment of mature toxin sequences from Cluster 1. The resulting model was visualized in Pymol. The superposition with Mu8.1 was generated in Pymol.

### Figures and Tables

**Table S1.** Available venom gland transcriptome datasets searched for Mu8.1- and con-ikot-ikot-like sequences

| *Conus species* | Accession numbers |
|---|---|
| *Conus epsicopatus* | DRX030966, SRR6983169, SAMD00029746 |
| *Conus raulsilvai* | SRR11807492 |
| *Conus infinitus* | SRR11807493 |
| *Conus antoniomonteiroi* | SRR11807494 |
| *Conus miruchae* | SRR11807495 |
| *Conus cuneolus* | SRR11807496 |
| *Conus boavistensis* | SRR11807497 |
| *Conus verdensis* | SRR11807498 |
| *Conus galeao* | SRR11807500 |
| *Conus maioensis* | SRR11807501, SRR11807499 |
| *Conus guanche* | SRR11807502 |
| *Conus grahami* | SRR11807507 |
| *Conus ventricosus* | SRR13740844 |
| *Conus bayani* | SRR13781584 |
| *Conus ebraeus* | SRR14407576, SRR14407590, SRR2609538 |
| *Conus mordeiraeo* | SRR14407578, SRR14407579 |
| *Conus regonae* | SRR14407580, SRR14407581 |
| *Conus fulgetrum* | SRR14407582 |
| *Conus abbreviatus* | SRR14407584, SRR14407585 |
| *Conus aristophanes* | SRR14407586, SRR14407587 |
| *Conus judaeus* | SRR14407589 |
| *Conus coronatus* | SRR14407591, SRR14407592, SRR2609545 |
| *Conus miliaris* | SRR1542424, SRR1542681, SRR1544117, SRR1544118, SRR1544119, SRR1544120, SRR1544137, SRR1544140, SRR1544142, SRR1544595, SRR1544597, SRR1544600, SRR1544622, SRR1544627, SRR1544690, SRR1544692, SRR1548185, SRR1548186, SRR1548187, SRR1548188, SRR1548189, SRR1548190 |
| *Conus betulinus* | SRR2124881 |
| *Conus ermineus* | SRR6983161, SRR6983162, SRR6983163, SRR6983164, SRR6983165, SRR6983166, SRR6983167, SRR6983168, SRR6983169 |
| *Conus magus* | SRR8195628, SRR9831255 |
| *Conus arenatus* | SRR2609544 |
| *Conus consors* | SRR1954994 |
| *Conus gloriamaris* | SRR5499408 |
| *Conus imperialis* | SRR12186674, SRR12186675, SRR12186676, SRR12186677, SRR12186678, SRR12186679, SRR2609542 |
| *Conus lividus* | SRR2609539 |
| *Conus marmoreus* | SRR8195632, SRR2609532 |
| *Conus quercinus* | SRR2609537, CNS0048932 |
| *Conus rattus* | SRR2609540 |
| *Conus rolani* | SRR16493597 |

| | |
|---|---|
| *Conus sponsalis* | SRR2609541 |
| *Conus striatus* | SRR8195630 |
| *Conus terebra* | SRR8195627 |
| *Conus textile* | SRR8195629 |
| *Conus tribblei* | SRR1799982 |
| *Conus varius* | SRR2609543 |
| *Conus virgo* | SRR8195631, SRR2608262 |
| *Conus geographus* | SRR503416, SRR503415, SRR503414, SRR503413 |
| *Conus trochulus* | SRR11807506 |
| *Conus reticulatus* | SRR11807504 |
| *Conus caracteristicus* | CNS0048931 |
| *Conus generalis* | CNS0048933 |

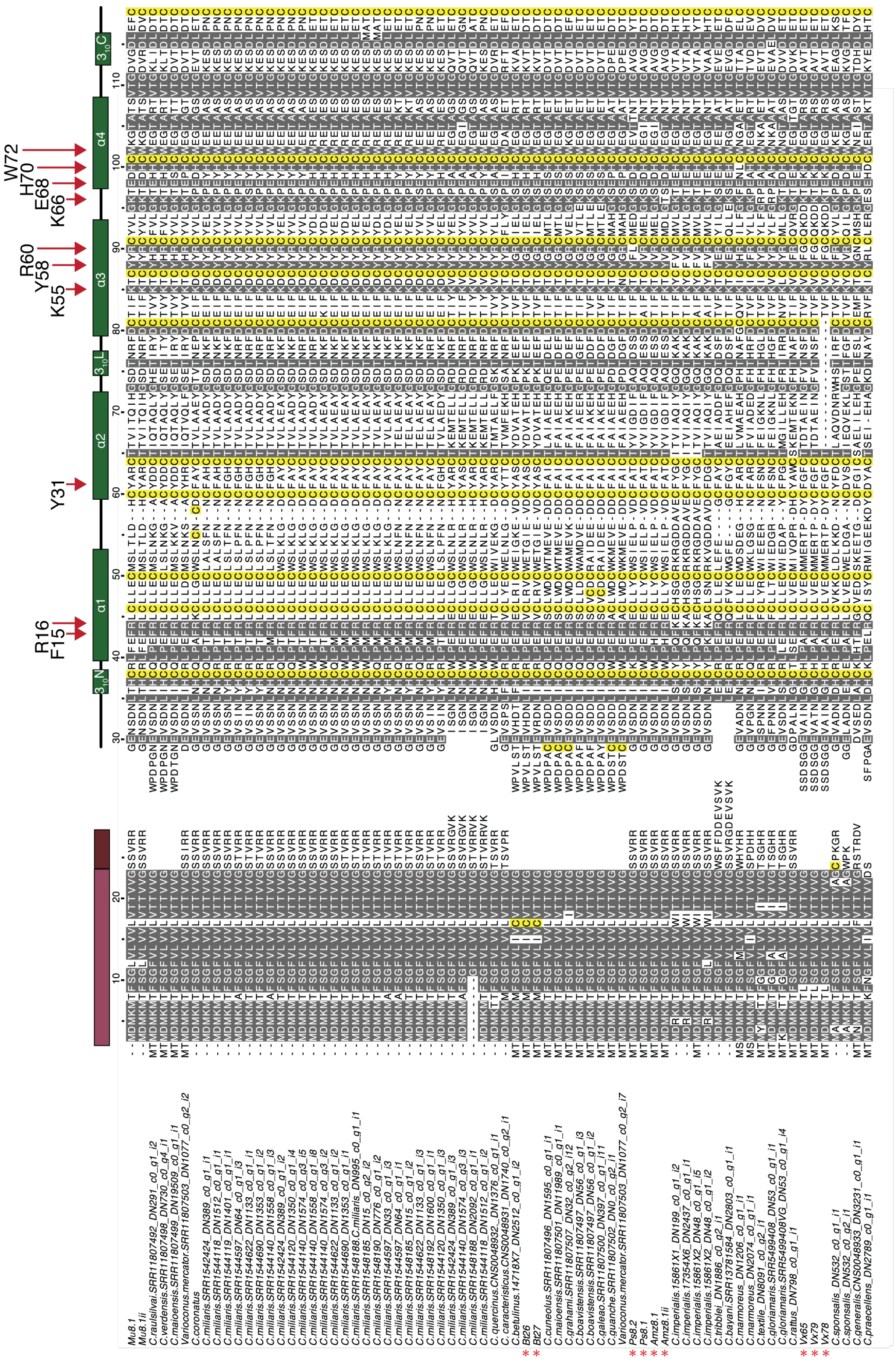

**Figure S1. Mu8.1 represents a new conotoxin superfamily**

Sequences harvested from venom gland transcriptomes available in the NCBI, DDBJ and CNGB repositories using the precursor sequence of Mu8.1 as query. Truncated sequences, duplicates and variants with only 1-2 amino acid residue differences were not included. The remaining sequences were used in a multiple sequence alignment carried out using the MAFFT version 7 multiple alignment online interface (11) and visualized in Jalview (12). For clarity, the signal and propeptide sequences are depicted with a space preceding the mature toxin sequences. Amino acid residues are shaded in gray according to a 90% identity threshold (all cysteine residues are shaded yellow regardless of conservation). Prepro-sequences are annotated with colored bars (positioned according to the Mu8.1 sequence) indicating the tripartite organization. Mauve: signal sequence, pink: propeptide region, green: mature conotoxin, here drawn to illustrate the α-helical structure as it corresponds to the Mu8.1 sequence. Red asterisks indicate those sequences previously annotated as con-ikot-ikot uncovered by pBLAST searching as described in the main text. Red arrows highlight amino acid residues referred to throughout the main text and green rectangles represent α-helices.

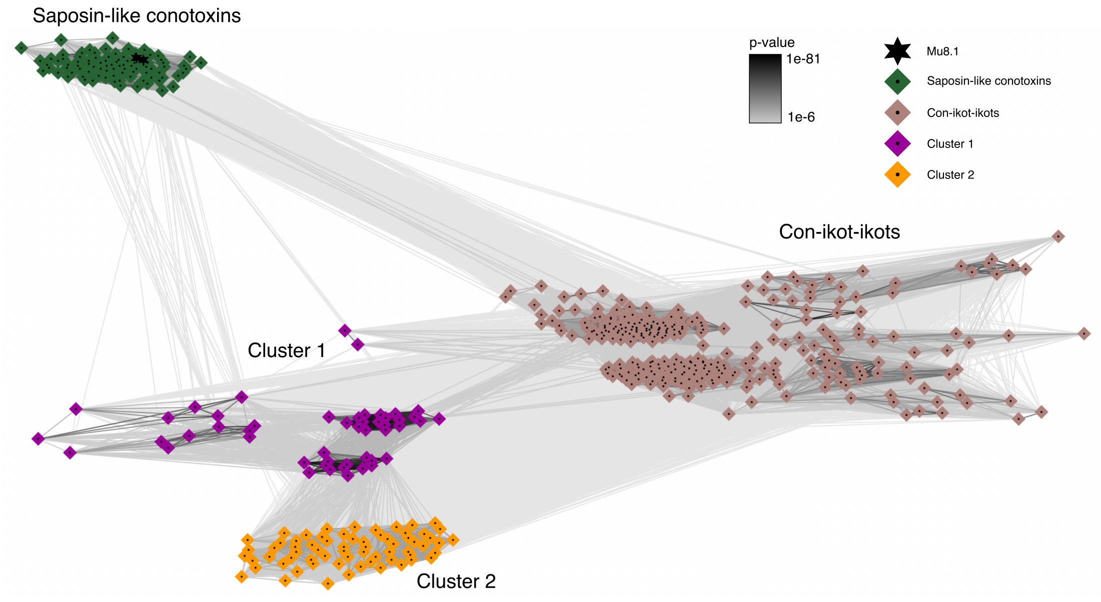

**Fig. S2.** BLOSUM62 cluster map of con-ikot-ikot-related conotoxin precursors. The nodes depict individual precursor amino acid sequences, and the edges correspond to the BLAST p-values < 1e-6 between the nodes. The four clusters are labeled by specific colors (Mu8.1 and Mu8.1ii are further highlighted by the star shapes), but the landmarks are also placed on the map.

```
Mu8.1 TRINITY_DN3539 gctctgtttcgcctactcgatcatacgccctagcagctgcagaccagcac 125
textile.TRINITY_DN80 gctgtgtctagtgtactcggccatacgccctagcagctgcagaccag-gc 149
18400X15.C.geographu ccctggttcagccagctctaccagacg-gcagacagctggggaccag-ac 125
SRR14407585.C.abbrev -------------------------------------tggggaccag-ac 12
SRR11807502.C.guanch tcccggcccgcactgctctgtcagatgcaaaaccacttggagaccaa-ac 61
varius_MP.TRINITY_DN gctctacccagtccgctctcccagtcgcccagcctctctcccatcgc-gc 53
SRR1548188.C.miliari ---------------------------------------------aa-ac 4
SRR14407585.C.abbrev -------------------------------------tggagaccaa-gc 12

Mu8.1 TRINITY_DN3539 acgtgaaaagaaagcattggaaacaagtgtggttatttcaagacacgcag 175
textile.TRINITY_DN80 acgtgaaaagaaagcattggaaacaagtgtggatatttcaagacatgcag 199
18400X15.C.geographu agacgtcaa-acagcatcgcagtcaggtgtggagatcccaaggcacccag 174
SRR14407585.C.abbrev agacgtcaa-acagcatcgtagtcaggtgtgcagatcccaagacacccag 61
SRR11807502.C.guanch ggacgttat-caagcacta-agtcgagtgaagagattccaagacaccgag 109
varius_MP.TRINITY_DN agacgttct--tcgcagtggaagcaagcggggtaattccagagt-ctcag 100
SRR1548188.C.miliari gggcgttga-caagcacta-agtcgagtcaagagattccaagacctcaag 52
SRR14407585.C.abbrev ggacgtcaa-caagcacta-agtcgagtgaagagattccaagacaccaag 60

Mu8.1 TRINITY_DN3539 a-acaaggaagcacaa---gtcatcgttcctg---aggcaataaccatgg 218
textile.TRINITY_DN80 a-acaaggaagcacag---gtcatcgttcccg---aggcaatgaccatgt 242
18400X15.C.geographu a-agaaggacacagaaga-gttatcgttcgta---atacaatggccatga 219
SRR14407585.C.abbrev g-agaaggaaacagaaga-gttatcgttcgtg---atacaatggccatga 106
SRR11807502.C.guanch acaaacggaagaagagga-gttgtcgttcttg---agaaaatggccacga 155
varius_MP.TRINITY_DN a-gcaaggaagcagaaaa-gttattgttcctg---agaaaatggctatga 145
SRR1548188.C.miliari acaaaaggacgacgaagaggttgtcattcttgaaaaaaaaatggccacga 102
SRR14407585.C.abbrev acagaaggacgaagag---gttgtcgttcttg---agaaaatggccacga 104

Mu8.1 TRINITY_DN3539 acatgaagatgacgtttagcgggttggtgctggttgtcttggtaaccact 268
textile.TRINITY_DN80 acatgaagacgacgttcggcgggtttgtgctggttgtcttggtaatcact 292
18400X15.C.geographu acatgtcggtgatgctcagcgcgtttgtaatggtcgtcgtgtcagccact 269
SRR14407585.C.abbrev acatgtggaagacgatcagcgtgtgtgttgtggttgtcatggcgaccact 156
SRR11807502.C.guanch acctttggatgacgctcggcatgctggtaatggttgtcatggcaaccgct 205
varius_MP.TRINITY_DN ccatacggatgacgatgaccgtggctgtactggctgtcatagcaaccacc 195
SRR1548188.C.miliari atctttggatgacgctcagcctgctggttctggttgtcatggcaaccact 152
SRR14407585.C.abbrev atctttggatgacgctcagcatgctggttatggttgtcatggcaaccgct 154
```

**Fig. S3.** Multiple sequence alignment of randomly selected (and Mu8.1) sequences representing the four toxin clusters. Only the 5' UTRs and beginning of ORFs are shown. The start codon is shown in yellow and columns with ≥ 75 % sequence identity are highlighted in black. The sequence names are colored according to match the clusters in Fig. S2.

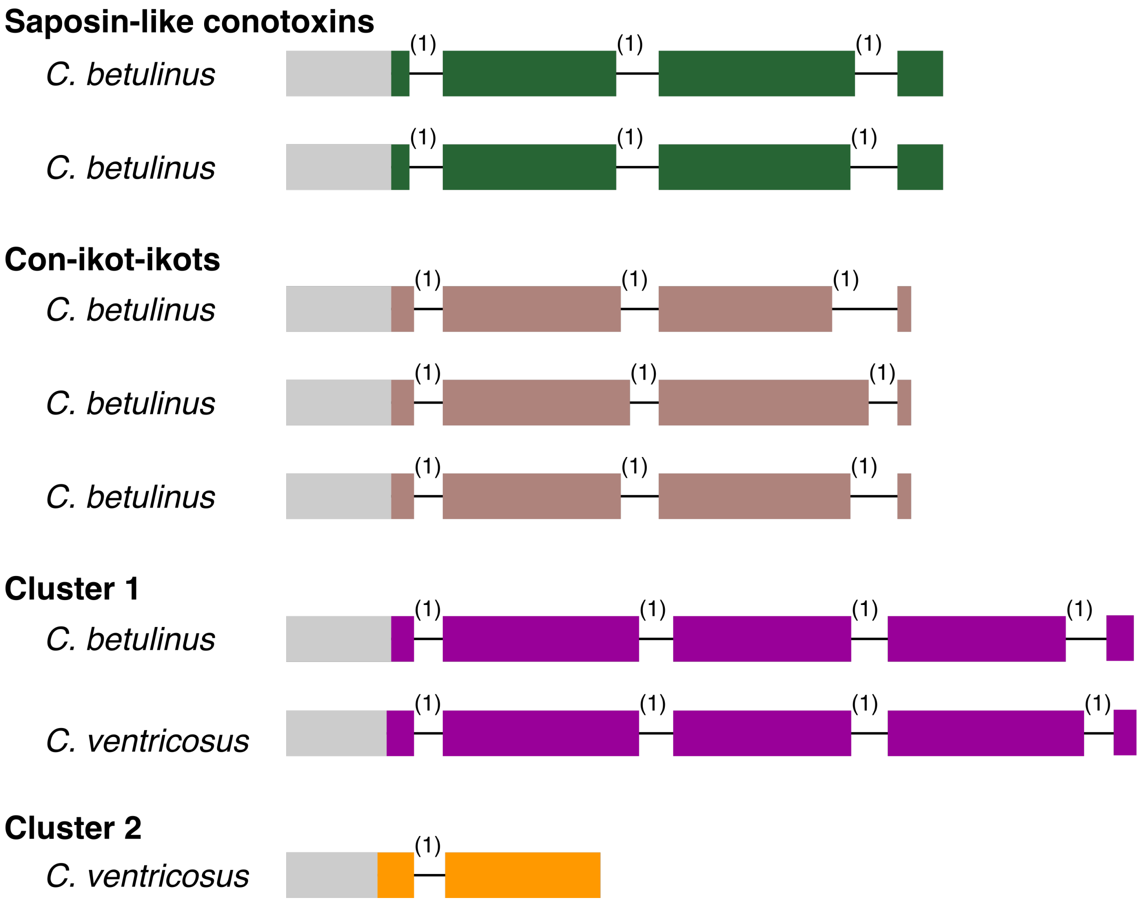

**Fig. S4.** Gene structure of *C. ventricous* and *C. betulinus* conotoxins from the four clusters in Fig. S2. The exons are represented by wide boxes proportional to the length of the sequences, whereas the introns are shown by thin interspaced segments (not proportional to sequence length) with their phases given above each intron. The predicted signal sequence is colored grey, and the remaining precursor colored to match Fig. S2.

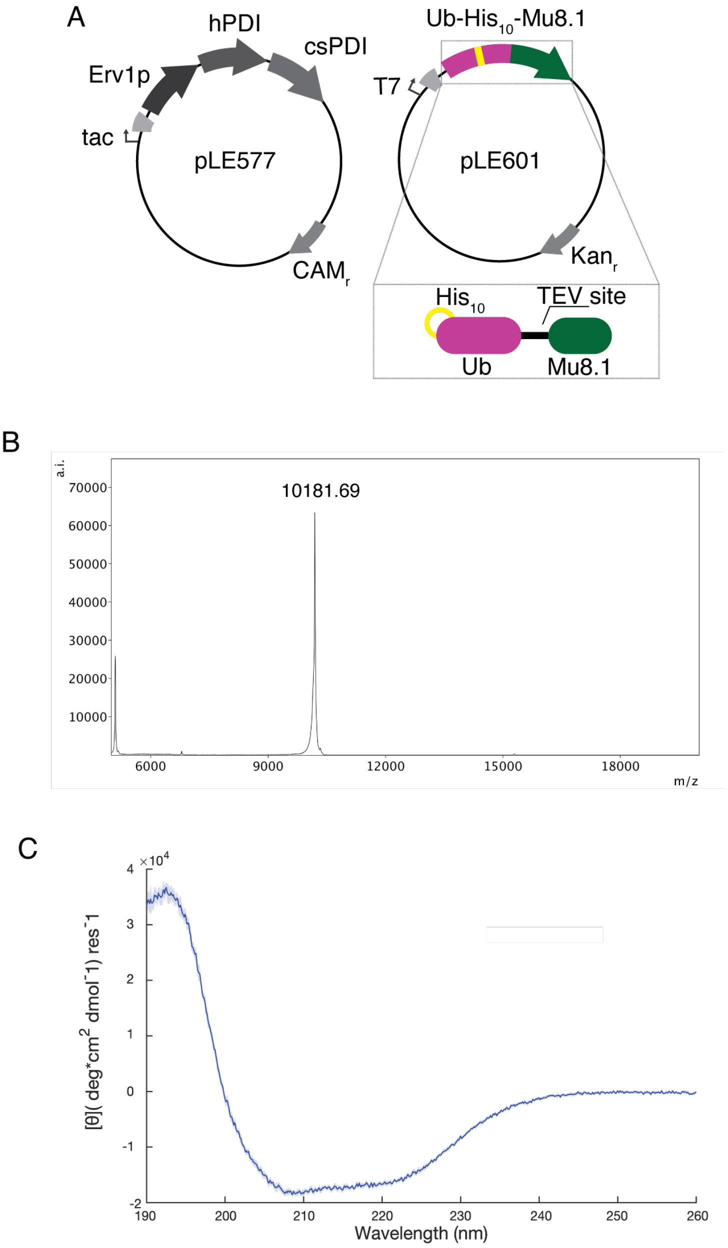

**Figure S5. A.** Schematic overview of the two plasmids of the csCyDisCo expression system (13) used for recombinant expression of Ub-His$_{10}$-Mu8.1. The csCyDisCo plasmid (pLE577) encodes for the three enzymes, Erv1p, human PDI (hPDI) and conotoxin-specific PDI (csPDI). pLE601 encodes the Ub-His$_{10}$-Mu8.1 fusion protein as shown in the light gray box. **B.** MALDI-TOF spectrum of non-reduced Mu8.1 showing a single peak with a mass of 10181.7 Da. The theoretical average mass of Mu8.1 is 10181.5 Da. **C.** CD spectrum of non-reduced Mu8.1 recorded at 25°C. The blue line represents an average of ten scans. Blue shading around the curve signifies the standard error of the mean of the ten recorded spectra.

A

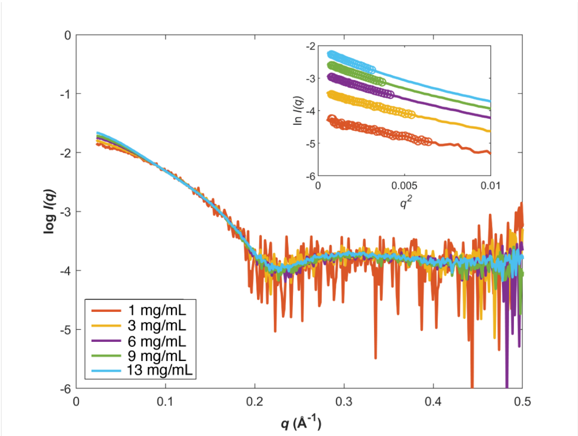

Table S2

| Sample | Rg, Guinier | Rg P(r) | I(0) Guinier | I(0) P(r) | Dmax | Mw Bayes | Mw Vc | Primus scale to 1 mg/mL sample | Est concentration from scale | Mw from scaled conc |
|---|---|---|---|---|---|---|---|---|---|---|
| 1mg/mL | 18.6 | 17.1 | 0.0145 | 0.0137 | 45 | 18.1 | 18.9 | 1 | 1.00 | 20 |
| 3 mg/mL | 20.4 | 20.6 | 0.051 | 0.0507 | 65 | 18.7 | 20.6 | 0.3231 | 3.10 | 22.8 |
| 6 mg/mL | 22.7 | 23.3 | 0.1214 | 0.1216 | 80 | 23.7 | 23.7 | 0.1584 | 6.31 | 26.7 |
| 9 mg/mL | 23.8 | 24.1 | 0.1786 | 0.1779 | 79 | 25.6 | 25.8 | 0.1175 | 8.51 | 29.1 |
| 13 mg/mL | 25.5 | 26.2 | 0.3143 | 0.315 | 89 | 28.9 | 28.7 | 0.0755 | 13.25 | 32.7 |

Table S3

| Sample | Conc mg/mL | Monomer % | Dimer % | Tetramer % | Octamer % | $\chi^2$ |
|---|---|---|---|---|---|---|
| 1 mg/mL | 1 | 0 | 85 | 15 | 1 | 0.94 |
| 3 mg/mL | 3.1 | 0 | 72 | 25 | 4 | 0.96 |
| 6 mg/mL | 6.3 | 0 | 57 | 37 | 6 | 1.03 |
| 9 mg/mL | 8.5 | 0 | 47 | 46 | 8 | 1.14 |
| 13 mg/mL | 13.2 | 0 | 34 | 55 | 12 | 1.15 |

**Figure S6. Mu8.1 exists primarily as a dimer in solution**

**A.** SAXS scattering profiles of increasing concentrations of Mu8.1 dissolved in 10 mM NaPi, pH 8, 150 NaCl. Inset: Guinier plots of scattering profiles, where straight lines were obtained by linear regression of the scattering profiles in the low q2 region. The Guinier region is highlighted with circles.

**Table S2. Data reduction and primary analysis of SAXS data**. Data extraction was performed using RAW. Rg guinier; radius of gyration (Rg) calculated from the slope of the Guinier plot (Panel A, inset); Rg P(r), radius of gyration obtained from pairwise distribution function; I(0)guinier, intensity at zero scattering angle extrapolated from Guinier plot; I(0) P(r), intensity at zero scattering obtained from pairwise distribution function; Dmax, maximum dimension; Mw Bayes, molecular weight from Bayesian Inference; Mw Vc, molecular weight from volume of correlation; Primus scale to 1 mg/mL sample, scaling obtained using the Primus program from the ATSAS package (14); Est concentration from scale, estimated concentration from Primus scale in mg/ml; Mw from scaled concentration (conc), molecular weight as a function of concentration.

**Table S3. Percentages of oligomeric species in solution at different concentrations of Mu8.1**. The percentages of oligomeric species from monomer to octamer were determined with Oligomer (15) from the ATSAS package (14).

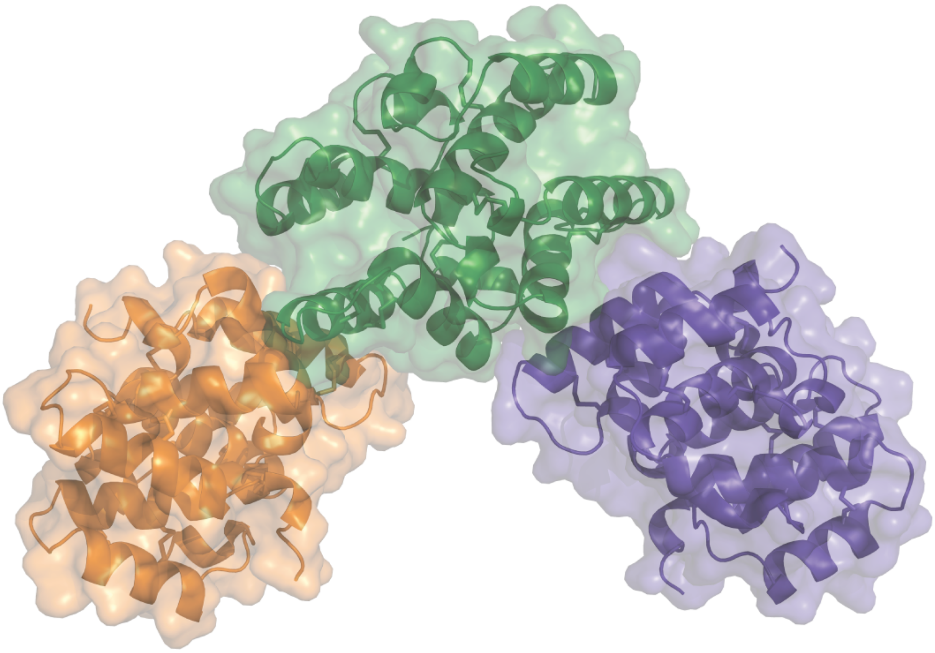

**Fig. S7. Molecules of the asymmetric unit of Mu8.1_59**

The asymmetric unit of Mu8.1_59 accommodates six molecules that form three equivalent dimers. The dimers are shown in orange, green, and dark purple.

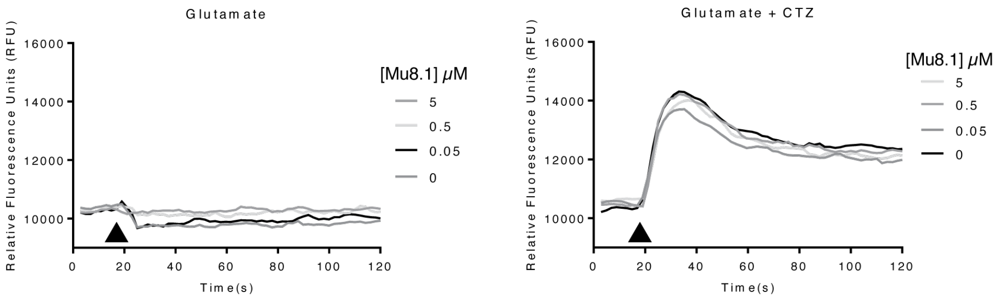

**Figure S8. Mu8.1 does not block desensitization of the GluA2 AMPA receptor**
Intracellular $Ca^{2+}$ imaging was used to determine a possible effect of Mu8.1 on the AMPA Receptor GluA2 as described in Materials and Methods. The experiment was performed in the absence (left) and presence (right) of cyclothiazide (CTZ), a positive allosteric modulator of AMPA receptors, known to block receptor desensitization and thus increase AMPA receptor current. Black arrows indicate the addition of saturating agonist solution (1 mM glutamate). The experiment was conducted at different concentrations of Mu8.1 as indicated. No effect of Mu8.1 treatment was observed.

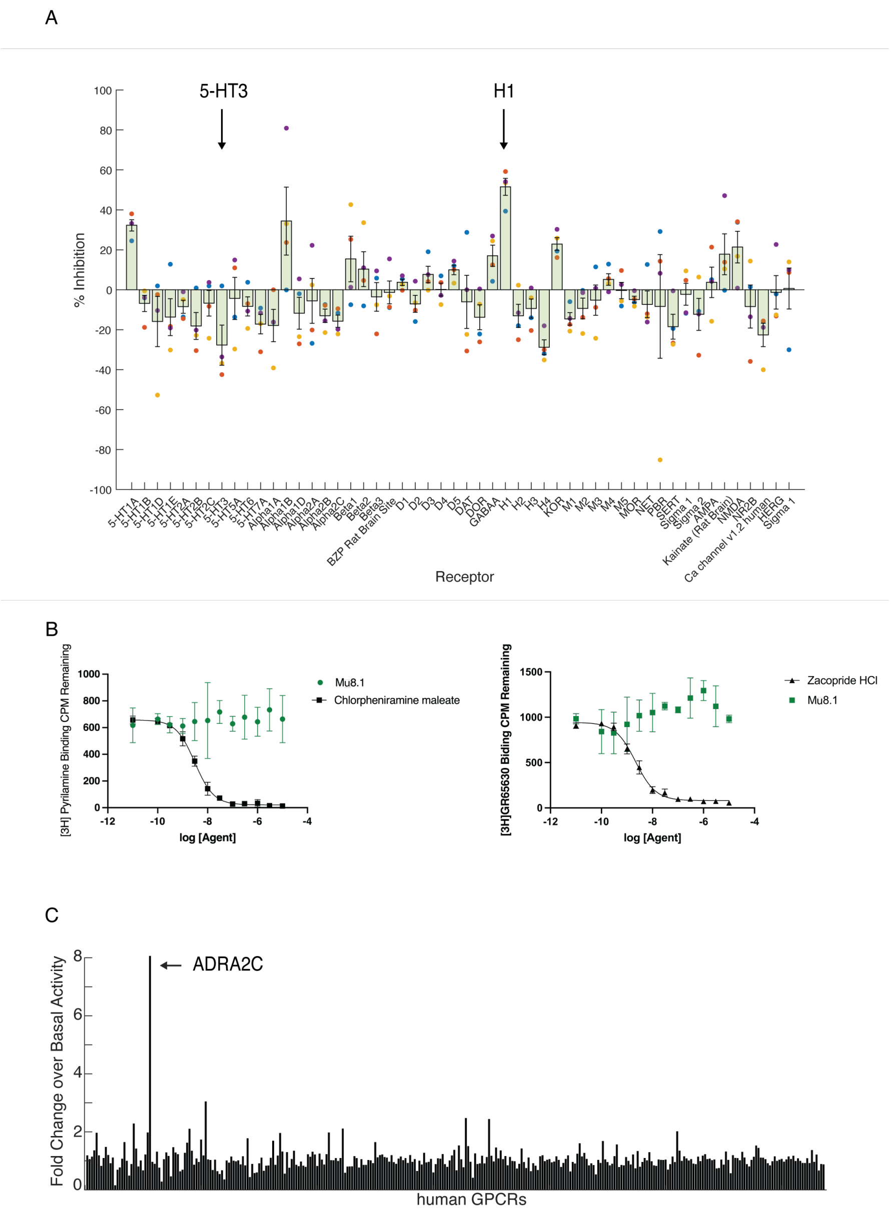

**Figure S9. Receptor and GPCRome screening of Mu8.1**

**A.** Primary radioligand binding assay of Mu8.1 (10 uM) at 52 receptors and ion channels (x-axis). Plotted values represent means (n=4) and error bars are ±SEM. Arrows show two receptors, 5-HT3 and histamine H1, with inhibition >50% that were selected for secondary testing. **B.** Secondary binding assay of the 5-HT3 (left) and H1 (right) receptors. Plotted values represent means (n=3) and error bars are ±SEM. The results demonstrate that the two initial hits were false positives. **C.** PRESTO-Tango screen of Mu8.1 (10 μM) at 318 human GPCRs (x-axis). Plotted values represent means ±SD. The arrow indicates potential hit (ADRAC2) that met the cut-off for further testing. However, a closer examination of the data showed a false positive due to an outlier.

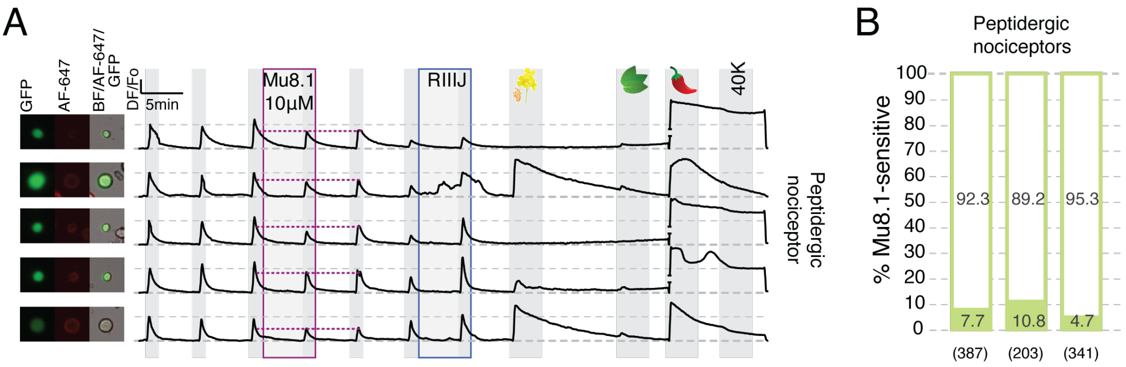

**Figure S10. Mu8.1 irreversibly inhibits $Ca^{2+}$ influx in a subpopulation of peptidergic nociceptors**

**A.** Examples of calcium traces from peptidergic nociceptors in which Mu8.1 inhibition of the peak was not reversed upon washout. Treatment with 10 µM Mu8.1 was scored as "irreversible" if the second $Ca^{2+}$ peak after Mu8.1 treatment was lower than the peak immediately before Mu8.1 treatment (see mauve dotted line). Each trace represents the calcium signal (ΔF/Fo) of the neuron pictured on the left (GFP-CGRP+: peptidergic nociceptors; Alexa Fluor 647-Isolectin B4+: non-peptidergic nociceptors, and bright-field). KCl depolarization pulses (25 mM) are indicated by light grey shading. A higher KCl pulse (40 mM) was used to elicit maximum calcium signal at the end of the experiment. Class-defining pharmacology: RIIIJ (1 µM, blue box); allyl isothiocyanate (AITC, 100 µM; mustard flower), menthol (400 µM; peppermint leaf), and capsaicin (300 nM; chili pepper). **B.** Peptidergic nociceptor populations from three independent experiments showing the percentage of neurons (within the bars) where Mu8.1 treatment was reversible (empty) or deemed irreversible (filled). The number of peptidergic nociceptive neurons recorded in each experiment is shown within parentheses.

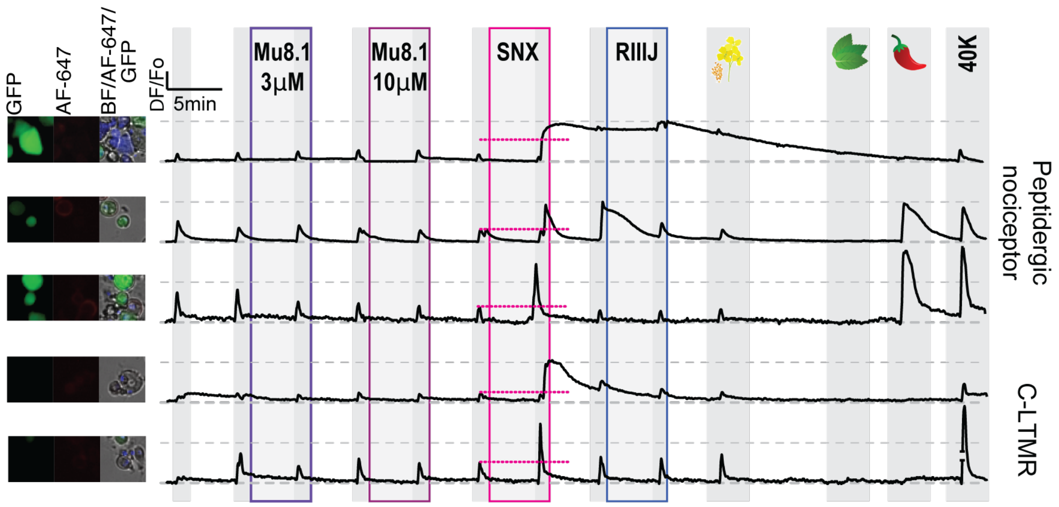

**Figure S11. SNX-482 amplifies intracellular $Ca^{2+}$ signal in subpopulations of sensory neurons**

Examples of calcium traces from peptidergic nociceptors and C-LTMRs in which SNX-482 amplified the KCl depolarization-induced $Ca^{2+}$ peak (see pink dotted line). Each trace represents the calcium signal (ΔF/Fo) of the neuron pictured on the left (GFP-CGRP+: peptidergic nociceptors; Alexa Fluor 647-Isolectin B4+: non-peptidergic nociceptors, and bright-field). KCl depolarization pulses (25 mM) are indicated by light grey shading. A higher KCl pulse (40 mM) was used to elicit maximum calcium signal at the end of the experiment. Class-defining pharmacology: RIIIJ (1 µM, blue box); allyl isothiocyanate (AITC, 100 µM; mustard flower), menthol (400 µM; peppermint leaf), and capsaicin (300 nM; chili pepper).

A

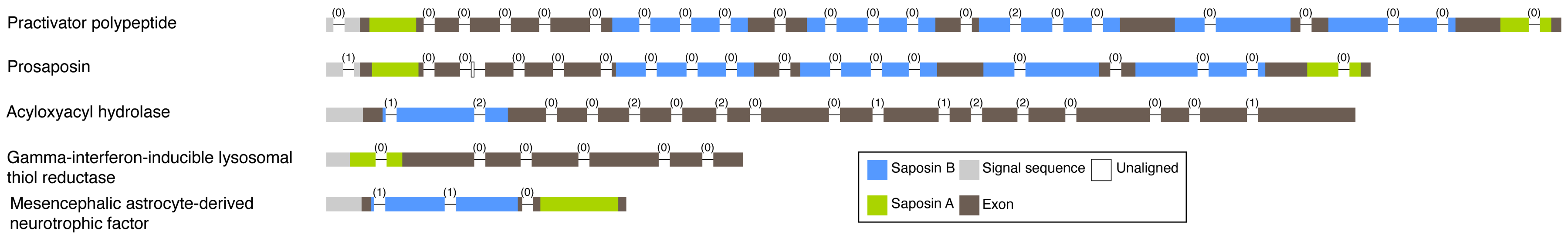

B

```
Mu8.1  GENSDNLTHCRLFEFRLCLLECMSLTLDHCYARCTTVITQIHGSDTNRFDCTIFKTC-------YYRCYVLGKTEDHCWKGTATSVTGDVG-------------------DLEFC-
MANF   RD-------CEV---------CISV-----LERFVNQTSEKDLKDLEKSEVKFEKFCAGLKGRDHKICYYLGAVKN-----APTRIVREMTKPLSFHMPPKKVCEKLKKHDAEICS
```

**Fig. S12. A.** Gene structure of saposin domain-containing proteins in *C. ventricosus*. The exons are represented by wide boxes proportional to the length of the sequences, whereas the introns are shown by thin interspaced segments (not proportional to sequence length) with their phases given above each intron. **B.** Sequence alignment of the saposin domains from Mu8.1 and *C. ventricosus* mesencephalic astrocyte-derived neurotrophic factor (MANF) revealing only very little sequence similarity.

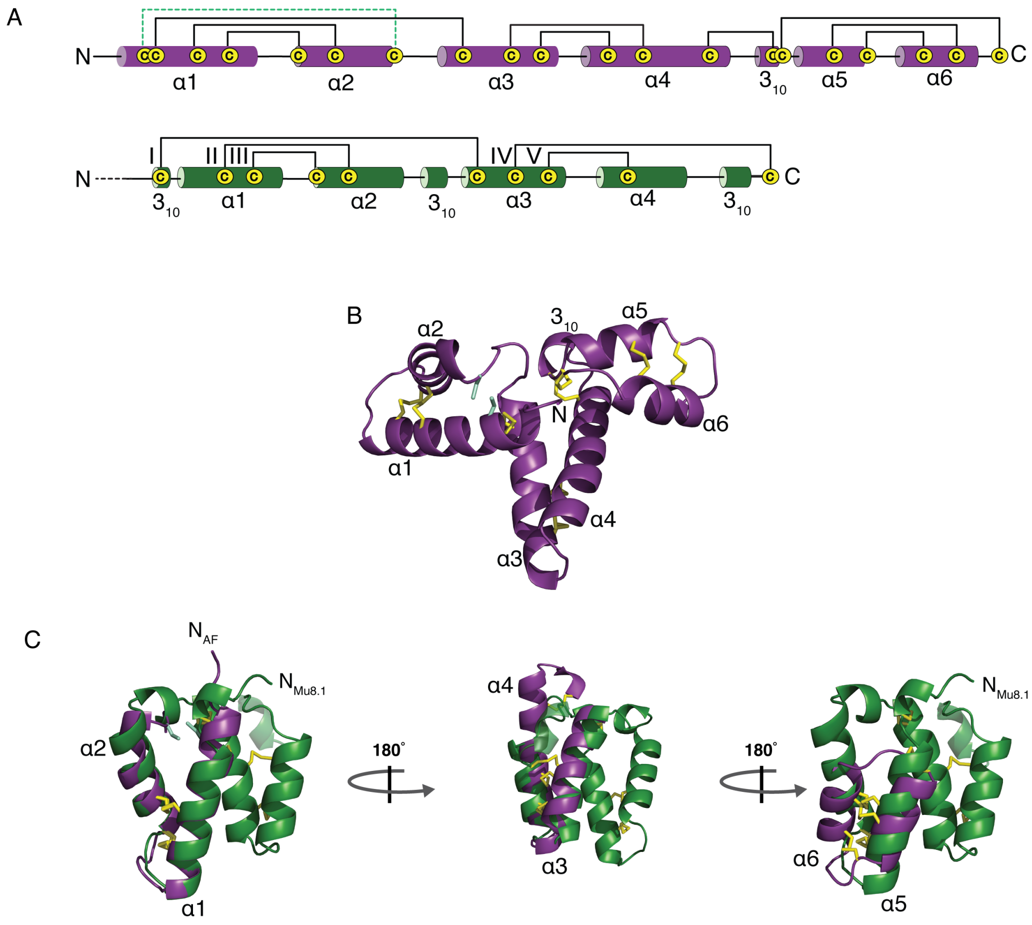

**Fig. S13. Toxins from Cluster 1 likely contain a saposin fold**

**A.** Graphic representations of AlphaFold-predicted secondary structure elements and disulfide bonds in a Cluster 1 conotoxin from *C. litteratus* (Supplementary file A and (8) (purple) and in Mu8.1 (green). Disulfide bonds are represented by brackets and α- and $3_{10}$-helices represented by cylinders (numbered by arabic numerals). The dotted pale green bracket represents a putative disulfide bond not predicted by AlphaFold. **B.** Cartoon representation of the structure predicted byAlphaFold for the Cluster 1 toxin. The predicted structure displays an α-helical protein with three leaf-like domains each consisting of a helix-turn-helix motif. Disulfide bonds are represented by yellow sticks and free cysteines are represented by pale green sticks. **C.** The three helix-turn-helix motifs individually overlaid with the Mu8.1 structure. Note the close structural similarity (and disulfide pattern) between the first four helices of the Cluster 1 protein and Mu8.1, apart from a predicted different orientation between the two helix-turn-helix motifs.

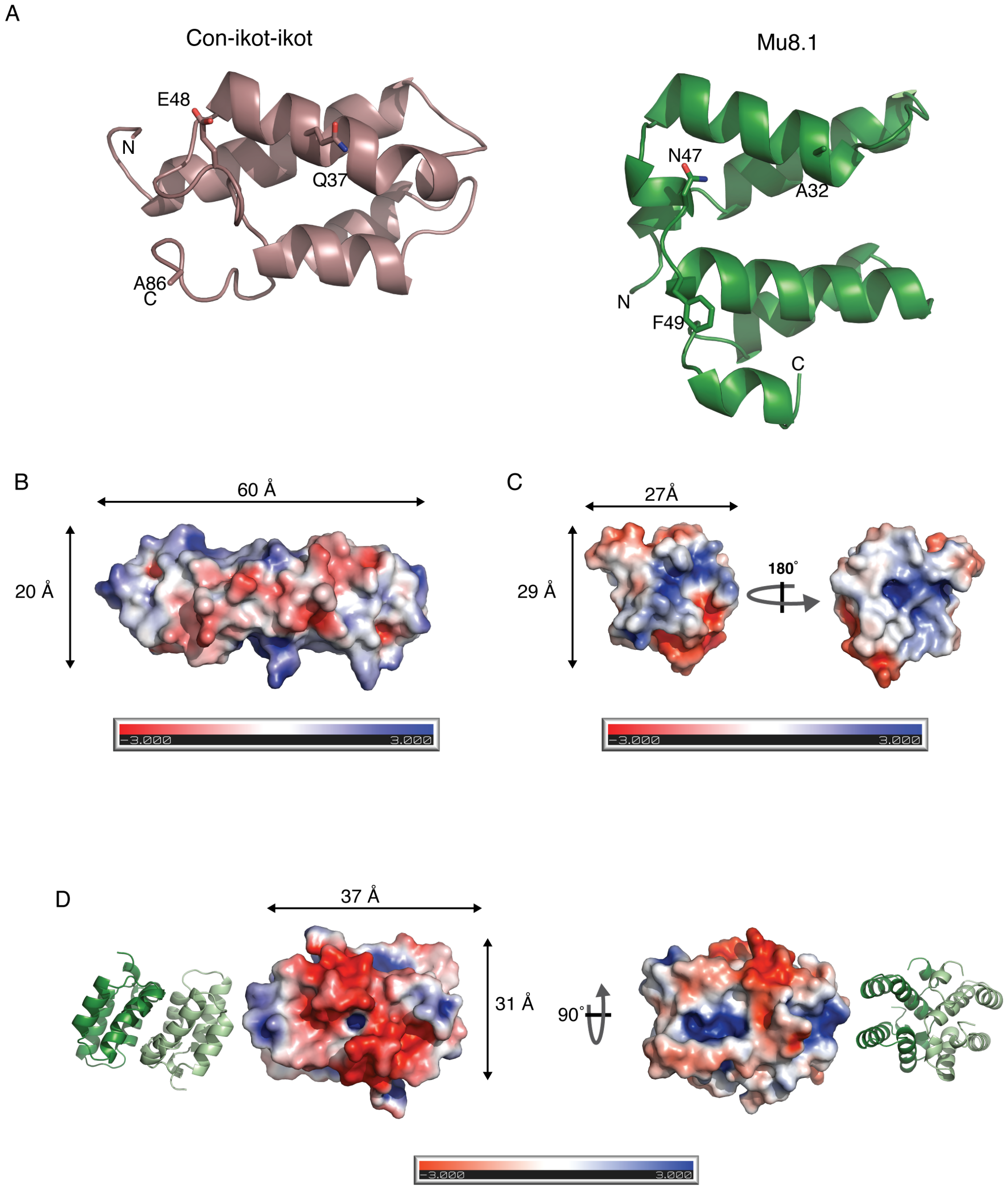

**Figure S14. Structure and sequence features explain Mu8.1's lack of activity at AMPA Receptor GluA2. A.** The GluA2 AMPA receptor-binding surface of con-ikot-ikot with residues important for binding (16) shown as stick models (left) and the corresponding surface and residues in the Mu8.1 protomer (right). **B.** Electrostatic surface representation (red: negative, blue: positive) of the con-ikot-ikot GluA2 AMPA receptor-binding surface (16). **C.** Electrostatic surface representation of the Mu8.1 protomer – left; outer surface of the dimer, right; surface of the dimer interface. **D.** Electrostatic surface representation of the Mu8.1 dimer showing a negatively charged patch (left). The cartoon representations next to each surface representation indicates orientation.

**Table S4**

| Ion Channel | Test Potential | [Mu8.1] | Inhibition | | | $IC_{50}$* |
|---|---|---|---|---|---|---|
| | (mV) | (μM) | (%) | | | (μM) |
| | | | Mean | SEM | n | |
| Cav2.1 | 0 | 10 | 26 | 4 | 2 | 28.5 |
| Cav2.2 | 10 | 10 | 19 | 7 | 3 | 42.6 |
| Cav3.1 | -20 | 10 | 20 | 1 | 3 | 40.0 |
| Cav3.2 | -20 | 10 | 23 | 3 | 3 | 32.5 |
| Cav3.3 | -20 | 10 | 19 | 3 | 2 | 42.6 |
| Kv1.1 | 20 | 10 | 19 | 5 | 5 | 42.6 |
| Kv1.2 | 20 | 10 | 7 | 4 | 3 | 132.9 |
| Kv1.3 | 20 | 10 | NB | | 6 | - |
| Kv4.3 | 20 | 10 | NB | | 5 | - |
| hERG | $20^{\#}$ | 30 | NB | | 3 | - |
| Nav1.2 | -10 | 30 | NB | | 4 | - |
| Nav1.4 | -10 | 30 | NB | | 3 | - |
| Nav1.7 | -10 | 30 | 9 | 3 | 4 | 303.3 |

**Table S4. Summary of activity determination of Mu8.1 against voltage-gated ion channels.** Automated patch clamp single concentration Mu8.1 screening of recombinantly expressed ion channels. Data for Cav2.3 is provided in Fig. 8. Mu8.1 concentrations, test pulse, and $IC_{50}$* estimates from single concentration experiments for each channel tested. $IC_{50}* = \{fc/(1-fc)\} \times [Mu8.1]$; $fc = I_{Mu8.1}/I_{Ctr}$; fc: fractional current; $I_{Mu8.1}$: current in the presence of Mu8.1; $I_{Ctr}$: current in the absence of toxin. NB = no block; SEM = standard error of the mean; n = number of experiments.
